# Splicing factor proline- and glutamine-rich (SFPQ) protein causes transcriptional repression of SNAIL to counteract TGF-β signaling

**DOI:** 10.64898/2025.12.01.691266

**Authors:** Niyati Pandya Thakkar, Hariharan Jayakumar, S Ramakrishnan, Gokula Narayanan, Srinjoy Chakraborty, Ritobrata Bhattacharyya, Yash Tushar Katakia, Suzanne L. Advani, Andrew Advani, Syamantak Majumder

## Abstract

TGF-β is known to regulate several embryonic and adult signaling pathways. Moreover, this signaling pathway regulates several cellular functions including differentiation, cell division, angiogenesis, hematopoiesis, and cell migration. However, studies suggest that an uncontrolled activation of TGF-β signaling may contribute to many human diseases. Therefore, counter-regulatory mechanism(s) to restrain abrupt TGF-β activation during cellular homeostasis is necessary to maintain an adequate balance of TGF-β downstream signaling. TGF-β through Smad complex activation causes transcriptional regulation of many transcription factors including Snail which act as an immediate-early response gene in TGF-β signaling. Herein, for the first time, we report that Splicing factor proline- and glutamine-rich (SFPQ), an RNA binding paraspeckles-associated protein works as a transcriptional repressor of Snail. We first confirmed a significant reduction in the expression level of SFPQ in the kidney glomeruli of rats that underwent subtotal nephrectomy. Endothelial cells (EC) treated with TGF-β exhibited loss of SFPQ protein level without altering its transcript level. Inhibition of proteasomal or autophagosome-lysosome pathway revealed ubiquitination-dependent proteasomal degradation of SFPQ upon TGF-β challenge. Prior to degradation, TGF-β treatment resulted in the cytosolic export of SFPQ thereby diminishing nuclear SFPQ level. Knockdown of SFPQ augmented TGF-β-dependent increase in Snail level while overexpression of SFPQ reversed TGF-β induced Snail expression. Although SFPQ exhibited association with many transcription factors including Smad2/3, Smad4, and N1-ICD which regulate Snail gene expression, TGF-β failed to alter the association of SFPQ with these transcription factors. Instead, through ChIP-qPCR analysis, we confirmed the enrichment of SFPQ in E-box promoter region and coding region proximal to TSS of the Snail gene. This study is the first to report SFPQ as a transcriptional repressor of Snail thereby regulating TGF-β signaling during cellular homeostasis.

## Introduction

Transforming Growth Factor β (TGF-β) is a key contributor of endothelial-to-mesenchymal transition (EndMT), governing the extent to which endothelial cells (ECs) lose their endothelial and acquire more mesenchymal characteristics. The capacity of ECs to undergo this transition is increasingly recognized as a source of pathological organ fibrosis [1]. TGF-β-dependent EndMT progresses with activation of both TGF-β I and II receptors, followed by canonical activation and translocation of Smad signaling complex (SMAD2/3 and SMAD4)[2]. In the nucleus, Smad complex induces the expression of key transcription factors like SNAIL, SLUG, TWIST, ZEB1, ZEB2, and FOXC2 that propels the cell to EndMT fate [3,4]. The ability to specifically modulate the EC response to TGF-β stimulation could offer unique opportunities to therapeutically offset the development of pathological fibrosis and maintain or restore endothelial function [5].

RNA-binding proteins (RBPs) control multiple aspects of mRNA biology, from splicing and export to stability and translation. They bind to specific regulatory regions in mRNA transcripts and exist as heterogeneous ribonucleoprotein (hnRNP) complexes that coordinate the expression of functionally related mRNAs. In addition to binding to RNA, RNA-binding proteins (RBPs) can also bind to DNA. The gene ontology analysis conducted by Hudson et al. revealed that DNA- and RNA-binding proteins (DRBPs) comprise approximately 2% of the human proteome [6]. Study by Imamura *et. al.,* showed that SFPQ binds to the promoter of the C-X-C motif chemokine ligand 8 (CXCL8)/interleukin 8 (IL8) genes, repressing their transcription. This transcriptional repressor function of SFPQ is regulated by the long non-coding RNA NEAT1. Mechanistically, NEAT1 derepresses CXCL8 expression by sequestering the transcriptional regulator SFPQ into paraspeckles, preventing its binding to the CXCL8 promoter [7,8]. Furthermore, SFPQ also binds to the promoter of the transcription factor FOS like antigen 1 (FOSL1) to activate its transcription in nasopharyngeal cancer cells. The long non-coding RNA LINC01503, which is regulated by the androgen receptor, has been shown to promote the recruitment of SFPQ to the FOSL1 promoter [9]. These multifunctional proteins play key roles in regulating crucial cellular processes, including transcription, mRNA processing, DNA replication, and DNA damage response (DDR). The broad regulatory roles of RBPs underscore their importance as central controllers of gene expression programs [6].

This highlights the critical regulatory functions of RBPs in gene expression programs. In addition to binding RNA, the ability of RBPs to bind DNA emphasizes their broad influence over cellular processes and disease pathogenesis, including neurodegenerative disorders, kidney diseases, diabetes, cardiovascular problems, and cancer[10]. However, the mechanisms of how paraspeckle-associated RNA binding proteins such as SFPQ control nuclear signaling pathways to regulate cellular homeostasis, specifically its role as a transcriptional repressor remains unclear. In this study, we provide strong evidence that SFPQ, a DNA- and RNA-binding protein acts as a transcriptional repressor of Snail through enrichment to Snail’s proximal promoter as well as coding region.

## Materials and Methods

### Cell culture

*In vitro* experiments were performed on multiple cells-including endothelial cell line and human embryonic kidney cells. EA.hy926 cells (immortalized human umbilical vein endothelial cells) (#CRL-2922, ATCC, USA) were cultured in Dulbecco’s modified Eagle’s medium (DMEM) (#AL006A, HiMedia Laboratories) supplemented with 10% fetal bovine serum (FBS, #RM1112, HiMedia Laboratories) and 1% penicillin/streptomycin (PS, #10378, Sigma-Aldrich, USA). Human Embryonic Kidney HEK-293 cell line was a kind gift from Prof. Uma Dubey (Department of Biological Sciences, Birla Institute of Technology and Science Pilani, Pilani Campus, India). HEK-293 were propagated in Minimum Essential Media Eagle (MEM) (#AT154; HiMedia Laboratories) supplemented with 5% FBS, and 1% PS. Cells were cultured at 37°C and in a humidified incubator with a 5% CO_2_ atmosphere.

### Treatment conditions

For cytokine treatment, EA.hy926 and HEK-293 cells were seeded at 5000 cells/cm2 and allowed to adhere for 24 hrs/overnight. Cells were treated with 10 ng/ml TGF-β1 (#100-21, PeproTech, NJ, USA) in 2% FBS supplemented DMEM/MEM and the medium was replaced post-48hrs. In general, short-term stimulations (0.5–12 hours) were applied to capture early signaling events, including post-translational modifications, protein interactions, subcellular localization, and transcriptional regulation, whereas extended treatments (24–72 hours) were used to assess downstream phenotypic and functional responses. To delineate the temporal dynamics of TGF-β–mediated regulation of SFPQ, multiple TGF-β treatment durations were systematically evaluated. Cells were collected for assessment at different time-intervals. For protein degradation studies, EA.hy926 cells were pre-incubated with proteasomal inhibitor and autophagy inhibitor, MG-132 (0.5 µM, #M7449, Sigma Aldrich, USA) and Bafilomycin A1 (#11038, Cayman chemicals, USA), respectively, for 4 h prior to cytokine treatment, cells were then collected at early and late time intervals. EC were pre-treated with SIS3 (1μM), a pharmacological inhibitor of TGF-β dependent Smad phosphorylation prior to inducing with TGF-β.

### siRNA silencing (inhibition studies) in cultured endothelial cells

Control siRNA (SignalSilence® Control siRNA #6568; Cell signaling Technology) was used at a concentration of 20-40 nM in combination with lipofectamine 2000 (#11668; Invitrogen, Thermo Fisher Scientific). Meanwhile, two different siRNA sequences were used to target SFPQ transcript (Supplementary Table 1), at the final concentration of 40-60nM. The siRNAs were incubated in Opti-MEM™ reduced serum medium (#31985; Gibco) for 4 h and subjected to the abovementioned treatment conditions. Scrambled siRNA was used as negative control. Cells were harvested for protein/RNA studies.

### Plasmid transfection

Exogenous overexpression of SFPQ gene were performed using pcDNA3.1 HA-SFPQ construct. HA-SFPQ was a gift from Rosalind Segal (Addgene plasmid # 166959; http://n2t.net/addgene:166959; RRID:Addgene_166959) [11]. Snail_pGL2 was a gift from Paul Wade (Addgene plasmid # 31694; http://n2t.net/addgene:31694; RRID:Addgene_31694) [12]. HEK-293 cells were seeded at density of 3000 cells/cm^2^ and were allowed to adhere overnight. Cells were incubated with plasmid (300ng/ml) and Lipofectamine 2000 (#11668, Invitrogen, Thermo Fisher Scientific) for 5 h. Post 24 h of incubation, cells were exposed to cytokine treatment as mentioned above and were collected for Immunoblotting and co-immunoprecipitation studies.

### Animal studies

Male Sprague Dawley rats (Charles River, Montreal, Quebec) aged eight weeks underwent sham or subtotal nephrectomy surgery as previously described [13]. The functional characteristics of these rats have been described previously [14]. Briefly, for subtotal nephrectomy surgeries, under isoflurane anesthesia, the right kidney was removed via subcapsular nephrectomy and infarction of two thirds of the left kidney was achieved by selective ligation of two out of three of the branches of the left renal artery. Sham surgery involved laparotomy and manipulation of both kidneys prior to wound closure. Rat kidney tissue was immersion-fixed in 10% neutral buffered formalin before routine processing and sectioning. All experimental procedures adhered to the guidelines of the Canadian Council on Animal Care and were approved by the St. Michael’s Hospital Animal Care Committee, Toronto, Ontario, Canada.

### Dual immunofluorescence of rat kidney tissue sections

Rat kidney sections (3 μm thick) from Sham and SNx rats were formalin-fixed and paraffin-embedded, followed by heat-induced epitope retrieval with citrate buffer. The sections were blocked (#X909, Protein Block; Dako), and incubated overnight at 4°C with JG12 antibody (1:100 dilutions; BMS1104, Thermo Scientific). On the subsequent day, after repeated washes with TBS-T, sections were incubated with secondary antibody Alexa Fluor 647 donkey anti-mouse (1:100 dilution; A31571, Thermo Fisher Scientific) for 2 h. Next, these sections were further incubated with SPFQ antibody (1:50 dilution; ab177149, Abcam) followed by the secondary antibody Alexa Fluor 488 donkey anti-rabbit (1:50 dilution; A21206, Thermo Scientific). DAPI was used at a concentration of 1:10,000. Scanning of stained slides were performed with a Zeiss Axioscan Z1 (Carl Zeiss Microscopy, Jena, Germany). Images were captured using a Zeiss LSM 700 widefield microscope and a 63X (1.4NA) objective controlled by Zen Black software (Carl Zeiss Canada, Toronto, Ontario, Canada). All images within a given experimental set were taken with identical acquisition settings. The antibodies used are specified in Supplementary Table 2. For measuring the relative level of SFPQ in the glomeruli of kidney tissues, we utilized the plugin of ImageJ for obtaining the level of green fluorescence in each glomerulus.

### Immunohistochemistry

Immunohistochemistry was performed with SFPQ antibody 1:50 (1:50 dilution; ab177149, Abcam). Incubation with phosphate buffered saline in place of the primary antibody served as the negative control. After incubation with the appropriate horseradish peroxidase conjugated Dako anti-rabbit secondary antibody at 1:10,000 dilution, sections were labeled with Liquid Diaminobenzidine and Substrate Chromogen (Dako North America Inc., Carpinteria, CA) before counterstaining in Mayer’s hematoxylin. Slides were scanned with the Aperio ScanScope System (Aperio Technologies Inc., Vista, CA).

### Immunofluorescence imaging and analysis of cultured cells

Glass coverslips were coated with 0.5% Gelatin for primary cells. Cells were cultured in 24-well plate (seeding density, 5000 cells/cm^2^). Cells were washed with PBS and fixed with 4% paraformaldehyde followed by permeabilization with 0.1% Triton X. Cells were incubated with blocking buffer (1% BSA, 0.1% TritonX-100) for 1 hour. The primary antibody was diluted in blocking buffer and incubated with coverslips overnight at 4. Coverslips were washed with PBST to remove unbound primary antibody and were incubated with Alexa fluor 488-conjugated anti-rabbit/mouse secondary antibody (1:4000) for 2 h. Cells were counterstained with DAPI (#D9542, Sigma-Aldrich) to visualize cellular nuclei. Fluorescence images were captured using a Zeiss LSM 880 Confocal microscope (Carl Zeiss). Fluorescence emission intensity was quantified using ImageJ software.

### Subcellular fractionation

Cells were harvested and washed thoroughly with PBS. The cell pellet was resuspended in ice-cold mild lysis buffer [NP-40 (0.1%, #MB143, Himedia Laboratories), Protease Inhibitor Cocktail (PIC,0.5%, #P8340, Sigma Aldrich), 1 mM Phenylmethylsulfonyl fluoride (PMSF, #TC706, Sigma Aldrich)]. The samples were then centrifuged for 10 minutes at 10,000 rpm and the nuclear fraction was obtained as pellet, whereas the supernatant was collected as the cytoplasmic fraction. Furthermore, the nuclear lysate was obtained by resuspending the nuclear pellet in harsh lysis buffer [1X RIPA buffer (#R0278, Sigma Aldrich), Glycerol (5%, #MB060, Himedia Laboratories), 1 mM PMSF #TC706, PIC (0.5%, #P8340, Sigma Aldrich). Cells were further lysed with sonication (15 kW, 2 cycles, 15sec each), followed by Immunoblotting. GAPDH and Histone H3 were used as controls for cytoplasmic and nuclear fraction, respectively.

### Immunoblotting studies

Media was removed and cells were briefly washed with sterile 1X PBS. Cells were incubated for 1 hour with RIPA buffer (#R0278, Sigma Aldrich) containing protease inhibitor (#P8340, Sigma Aldrich) for lysis on ice. Cells were scraped and collected, followed by repeated cycles of sonication (15 kW, 2 cycles of 15s each). Cell debris and whole cell lysate were separated by centrifugation (10,000g for 10 min). Protein concentration was estimated with Bradford assay (Absorbance at 595 nm). Protein samples were processed for SDS polyacrylamide gel electrophoresis, with 10-250 KDa pre-stained protein ladder (#PG-PMT2922, Genetix) as a molecular weight reference. Proteins were transferred onto a PVDF (*Polyvinylidene fluoride)* membrane at 15 V, 2.5 A for 30 minutes. The membrane was blocked with 3% Skimmed milk buffer (#GRM1254, Himedia Laboratories) for 1 hour. Membrane was probed overnight at 4°C by incubation using different monoclonal human/rat specific primary antibodies as specified in Supplementary Table 2.

### Co-immunoprecipitation studies

Cells were harvested and washed thoroughly with PBS. The cell pellet was resuspended in ice-cold mild lysis buffer [NP-40 (0.1%, #MB143, Himedia Laboratories), PIC (0.5%, #P8340, Sigma Aldrich), 1 mM PMSF #TC706]. The samples were then centrifuged for 10 min at 10,000 rpm and the nuclear fraction was obtained as pellet. Initial amount of 1 mg of nuclear lysate was used for pulldown with antibody for SFPQ (Sigma). Antibodies specific to SMAD4 (Abclonal), NICD (Abclonal), SMAD2/3 complex (Abclonal) were used at dilution of 1:1000. Resulting bands were visualized by Chemi-Doc using the Clarity™ (#1705061, Bio-Rad Laboratories) or Clarity™ Max (#1705062, Bio-Rad Laboratories) Western Blotting ECL Substrates and analyzed by expression levels relative to 5% input obtained using ImageJ software.

### RNA Isolation and Reverse Transcriptase-Quantitative Polymerase Chain Reaction

Cells were cultured in 24-well plate (seeding density, 5000 cells/cm^2^). Following respective treatment, cells were harvested for RNA extraction according to manufacturer’s protocol (#15596, TRIzol™ Reagent, Life Technologies, Thermo Fisher Scientific). RNA pellets were resuspended in sterile nuclease-free water and quantified on BioTek Synergy H1 instrument. Isolated RNAs were pre-treated with DNase to remove DNA contamination. Initial amount of RNA (1 μg) was used for the cDNA preparation using iScript™ cDNA Synthesis Kit (#1708891, Bio-Rad Laboratories, Hercules, CA, USA). Real-time PCR was performed using iTaq™ Universal SYBR^®^ Green Supermix (#1725124, Bio-Rad Laboratories), with a total reaction volume of 10 μL. Analysis was carried out by calculating delta-delta Ct. GAPDH was used as the housekeeping gene. Primer sequences to analyze the transcript level of different genes are mentioned in the Supplementary Table 3.

### Protein-Chromatin binding studies

The SFPQ protein binding on Chromatin was assessed using CUT&RUN assay (#86652, Cell Signalling Technologies), as per the manufacturer’s protocol. HEK-293 cells were transfected with HA-SFPQ construct and incubated for 24 h. EA.hy926 (5×10^6^) and HEK-293 (4×10^6^) cells were harvested, washed, and crosslinked with 1% formaldehyde for 2 min at room temperature, neutralized with 125 mM glycine for 5min and washed with ice-cold PBS. Cells were bound to activated Concanavalin A coated magnetic beads and permeabilized. The bead–cell complex was incubated overnight with the SFPQ antibody at 4°C. Cells were washed three times and resuspended in 100 μl pAG/MNase and incubated for 1 hour at RT. Input chromatin was fragmented with probe sonicator for 40 cycles (30 sec Pulse, 20 sec OFF) on ice. Chromatin was eluted using Silica Spin columns. DNA eluent was subjected to real-time PCR assay using iTaq™ Universal SYBR® Green Supermix (#1725124, Bio-Rad Laboratories). The chromatin occupancy was calculated with the formula: chromatin occupancy= 2 ^ [△Ct of test site (ChIP-input)-△Ct of control site (ChIP-input)]. Primer sequences targeting multiple region of Snail promoter for qPCR are as mentioned in the Supplementary Table 4.

### Promoter Reporter Assay

HEK293 cells were transfected with a luciferase reporter under the control of the *SNAIL* promoter, Snail_pGL2. Cells were incubated either co-transfected with pcDNA3.1 HA-SFPQ construct or SFPQ siRNA followed by treating the cells with TGF-β for 24 h before determination of luciferase activity with a reporter assay system (#E1500, Promega, Madison, MI).

### Statistics

All the values are expressed as the mean ± SD. All analysis data in bar graphs are presented as relative to control treatment conditions. Statistical significance was determined by one-way ANOVA and a two-tailed Student’s t-test for comparisons between two groups (or a Mann– Whitney U test for nonparametric data). Statistical analyses were performed using GraphPad Prism software. A p-value of less than .05 was considered statistically significant.

## Results

### TGF-β caused reduction in SFPQ protein level and concurrent induction of Snail gene expression in endothelial cells without altering the quantity of other paraspeckles associated proteins

TGF-β is reported to induce EndMT both *in vitro* and *in vivo* conditions [15]. To evaluate the protein abundance of SFPQ, we stained kidney tissues of the rats that underwent SNx surgery. We analyzed the expression of SFPQ in this model as activation of TGF-β signaling and associated kidney fibrosis are common in such model [16]. In Sham animals, many glomerular cells including glomerular EC (red, as counterstained using JG12) showed robust expression of SFPQ which was primarily localized in the nucleus (Figure 1A). In contrast, glomerular EC in SNx rats exhibited significant reduction in nuclear SFPQ level (Figure 1A-B, C).

**Figure 1.**
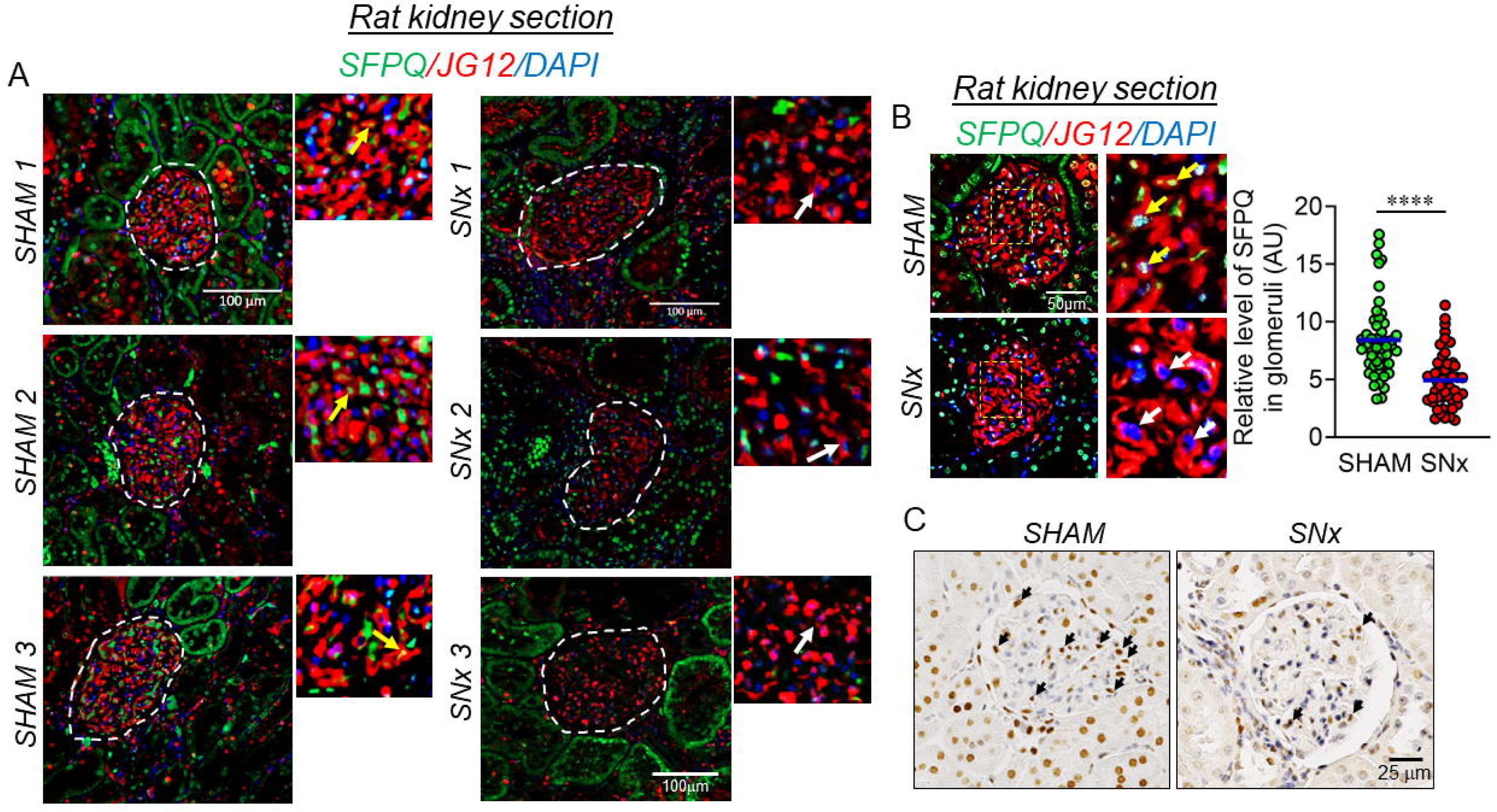
Diminished levels of SFPQ protein in glomeruli of the kidney tissue sections in subtotal nephrectomy (SNx) rats. (A-B) Immunohistochemistry of tissue sections from sham and sub-nephrectomy rats stained for SFPQ (Green, n=3) and JG12 (Red, n=3), DAPI staining to visualize the nucleus is shown in blue. Fluorescence intensity AU values per individual glomerulus are indicated together with the mean. Total number of glomeruli analyzed, n= 50. Yellow arrow heads show the presence of SFPQ (green) stained nucleus (blue) of the endothelial cells (JG12 positive cells, stained red) in the glomeruli of SHAM animals while white arrow heads indicate the absence of SFPQ (green) stained nucleus (blue) of the endothelial cells (JG12 positive cells, stained red) in the glomeruli of SNx animals. (C) Representative photomicrographs of SFPQ immunostained (developed with 3, 3’-diaminobenzidine) kidney sections from sham, and SNx rats. Black arrow heads show the presence of SFPQ stained nucleus in the glomeruli. Values represent the mean ± SD. **** p < 0.0001 by unpaired t test.

To confirm such diminished level of SFPQ *in vitro*, we treated cultured EA.hy926 cells with 10 ng/mL TGF-β at different time points. We first confirmed the activation of TGF-β dependent activation of Smad signaling by analyzing the level of p-Smad2/3 and confirmed increase in p-Smad2/3 in EC exposed to TGF-β for different time points (Figure S1A-B). Immunoblot analysis showed loss of SFPQ protein as early as 4 h, which remained low until 12 h post TGF-β challenge (Figure 2A). We also assessed the level of SFPQ, and Snail in EC treated with TGF-β for a different time point including 0.5, 1, 2, 4, and 8 h. In doing so, we observed that a reduction in SFPQ protein level was detected in EC 4 h post TGF-β treatment (Figure S1C). In contrast, Snail protein level enhanced within 0.5 h of TGF-β exposure (Figure S1C). We also detected time dependent increase in *SNAIL* transcript level in EC exposed to TGF-β (Figure S1D).

**Figure 2.**
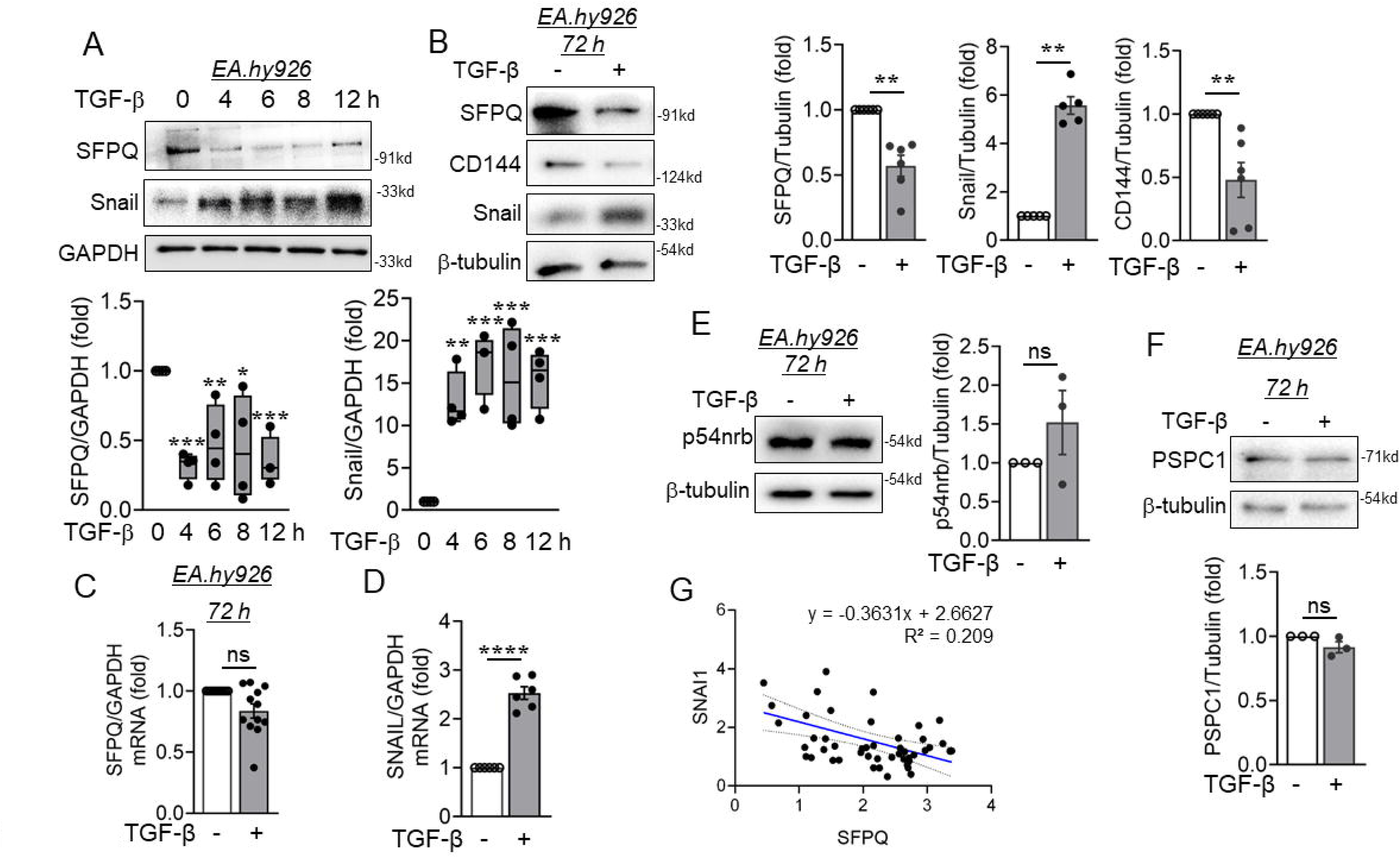
Endothelial cells subjected to TGF-β, showed enhanced expression of mesenchyme-specific proteins and reduced expression of SFPQ protein. (A) EA.hy926 cells were treated with TGF-β (10 ng/ml) and collected at different time intervals followed by immunoblotting for SFPQ and Snail protein (A, n=4). (B) Immunoblot analysis of EA.hy926 cell lysates collected from cells treated with cytokine TGF-β (10 ng/ml) for 72 h and probed for SFPQ (B, n = 6), CD144 (B, n = 6), and Snail (5, n = 6) along with their respective densitometry quantification analysis. (C-D) Transcript-level expression of SFPQ (C, n = 12) and SNAIL (D, n = 6) measured through RT-qPCR technique in EA.hy926 cells treated with TGF-β (10 ng/ml) for 72 h. All analyzed data were normalized to the control treatment condition. (E-F) Immunoblot analysis of EA.hy926 cell lysates collected from cells treated with cytokine TGF-β (10 ng/ml) for 72 h and probed for p54nrb (E, n = 3), and PSPC1 (F, n = 3) along with their respective densitometry quantification analysis. (G) correlation analysis between SFPQ and SNAIL mRNA expression levels using publicly available microarray data (GEO accession number: GSE66494) derived from kidney tissue samples of patients with chronic kidney disease (CKD, n = 48) Values represent the mean ± SD. * p < 0.05, ** p < 0.01, ***p < 0.001, and ****p < 0.0001 by unpaired t test or one way ANOVA.

Interestingly, we observed differences in expression level of *SNAIL* gene transcript and protein level when EC exposed to TGF-β for different time points (Figure S1C-D). Such data reflects post-transcriptional and post-translational regulatory mechanisms that are well-documented in the context of Snail gene expression. While Snail mRNA levels typically respond rapidly to TGF-β signaling, the translation of Snail protein, as well as its stability, is subject to complex regulation. Several studies have shown that Snail protein stability is tightly controlled by post-translational modifications such as phosphorylation [17], ubiquitination [18,19], and interactions with specific cofactors [20], which can delay or prolong its presence relative to mRNA expression. For instance, in most cells, Snail is a short-lived protein (with a half-life of about 25 min) and is rapidly ubiquitinated and degraded by the 26S proteasome [18]. Moreover, Snail1 is ubiquitinated by seventeen different E3 ubiquitin ligases, most of the Cullin-RING type. The Lys residues modified by these ubiquitin ligases have been identified only in a few cases, but they seem to be preferentially located in the SRD and NES subdomains [21]. Thus, the temporal disconnect between mRNA and protein levels likely reflects a combination of transcriptional activation, delayed translation, and altered post-translational stabilization of Snail protein in response to TGF-β. EC treated with TGF-β for 72 h also displayed restricted expression of SFPQ (Figure 2B). Besides, EC presenting diminished levels of SFPQ at different time points post exposure to TGF-β manifested elevated level of mesenchymal marker-Snail (Figure 2A-B). Reduced expression of endothelial markers-VE Cadherin was also observed in EC upon 72 h of TGF-β treatment (Figure 2B). However, when we examined the transcript level expression of SFPQ, it remained unchanged in EC exposed to TGF-β for 72 h (Figure 2C), while the transcript levels of Snail remained significantly upregulated in such experimental condition (Figure 2D). Interestingly, except SFPQ, other members of DBHS (Drosophila behavior, human splicing) class of proteins i.e. PSPC1 and p54nrb did not display any alterations in protein expression upon TGF-β stimulation (Figure 2E-F). We also performed a correlation analysis of mRNA levels of SFPQ and Snail using a GEO dataset (GEO accession number: GSE66494) published in a previous study [22]. This study reports the microarray analysis of renal biopsy specimens from 48 CKD patients in order to identify gene expression differences associated with CKD [22]. We observed a significant negative correlation (r = –0.4572) between Snail and SFPQ mRNA expression levels, consistent with our current findings demonstrating reduced SFPQ protein abundance under TGF-β treatment accompanied by increased Snail mRNA expression. These observations collectively underscore the clinical relevance of the inverse relationship between SFPQ and Snail (Figure 2G).

We next questioned whether TGF-β dependent activation of Smad signaling plays any role in governing SFPQ protein level in TGF-β treatment condition. We therefore used pharmacological inhibitor of Smad3 phosphorylation [23] and assessed the level of SFPQ upon TGF-β stimulation. We observed that pharmacological inhibition of Smad phosphorylation reversed TGF-β dependent reduction in SFPQ protein level (Figure S1E) indicating Smad activation plays crucial role in TGF-β mediated regulation of SFPQ protein level.

### TGF-β depleted SFPQ levels via proteasomal degradation pathway

Since no changes were detected in the transcript levels of SFPQ, and SFPQ protein loss occurred as early as 4 h after TGF-β treatment, we hypothesized that its degradation occurs post-translationally and began investigating the precise mechanisms involved in this process. Before analyzing the degradation pathway involved in TGF-β dependent SFPQ degradation, we first quantified the ubiquitination of SFPQ protein upon TGF-β challenge. Through immunoprecipitation and immunoblot analysis, we confirmed that SFPQ ubiquitination was significantly elevated upon TGF-β stimulation (Figure 3A). To identify the primary pathway responsible for SFPQ degradation upon TGF-β challenge, we inhibited major pathways i.e. ubiquitin-proteasomal and autophagosome-lysosomal mediated protein degradation machineries and further analyzed the levels of SFPQ protein using total cell lysates and immunoblot analysis. As noted earlier, we observed a significant increase in SFPQ ubiquitination upon TGF-β exposure. However, upon inhibition of ubiquitin-proteasomal protein degradation pathway with small molecule inhibitor MG-132 at early intervals, we detected observed a reversal in the SFPQ protein level in TGF-β stimulated EC (Figure 3B). Such reversal in SFPQ protein level in MG-132 exposed and TGF-β treated cells suggest a robust proteosomal pathway dependent degradation of SFPQ in EC upon TGF-β challenge. Notably, the accumulation of SFPQ in MG-132 and TGF-β co-treated condition exceeded that seen with MG132 treatment alone, suggesting a possible compensatory upregulation of SFPQ expression by cells in response to TGF-β stimulation. EC pre-treated with Bafilomycin A1 followed by TGF-β challenge was unable to reverse the protein abundance of SFPQ (Figure 3C). Inhibition of autophagic flux by Bafilomycin A1 was confirmed by detecting the level of p62 and LC3B (Figure 3C). Furthermore, we assessed K48-linked ubiquitination of SFPQ, a modification predominantly associated with proteasomal degradation, and found that TGF-β treatment increased the level of K48-linked ubiquitin conjugation on SFPQ (Figure 3D). These results indicate TGF-β regulates SFPQ abundance post-translationally via. proteasomal-dependent degradation. We also assessed the level of E3 ubiquitin ligases that may be enhanced in TGF-β challenged EC. In so doing, we observed that E3 ubiquitin ligases such as *RNF11*, *SMURF1*, and *NEDD4L* are significantly enhanced in EC exposed to TGF-β for 24 h (Figure S2A). In contrast, TGF-β did not alter the level of *SMURF2*, and *CYCLO* (Figure S2A).

**Figure 3.**
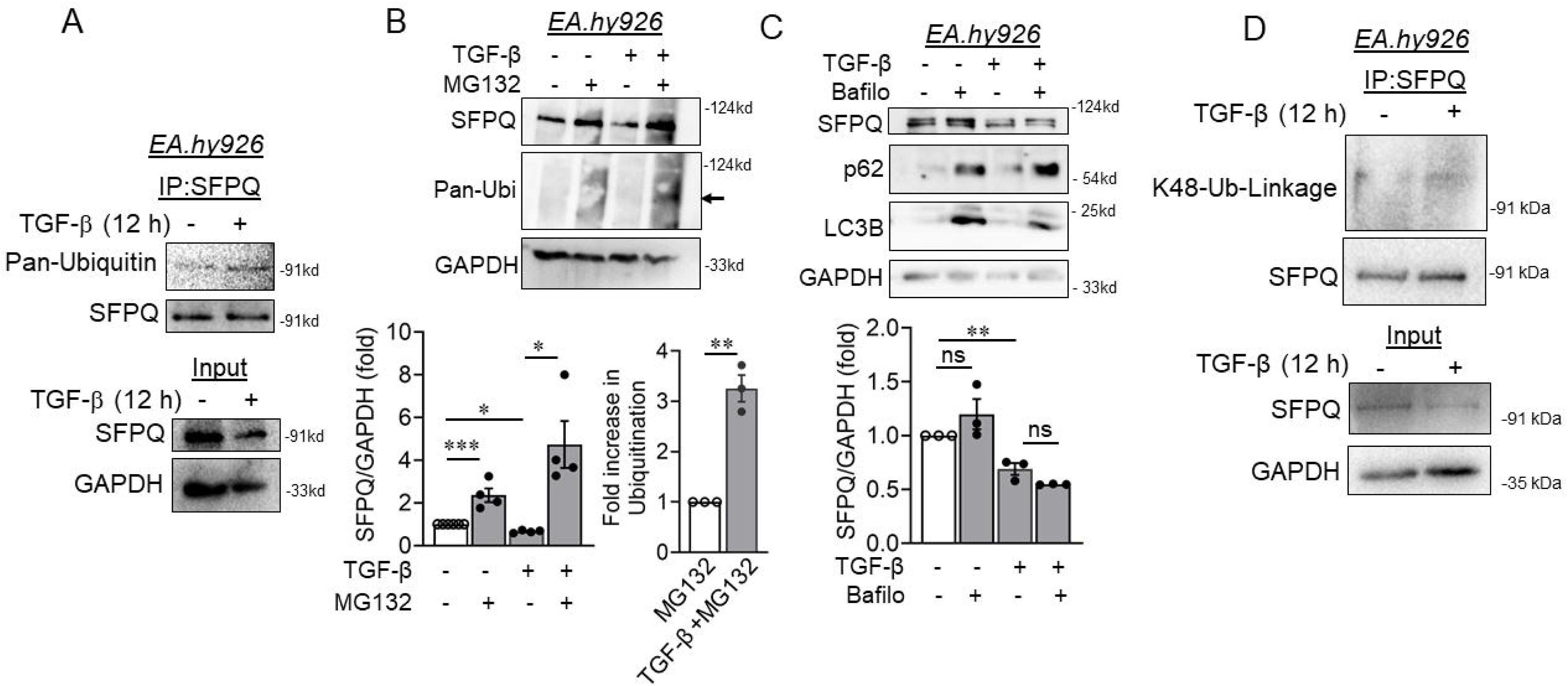
TGF-β induction caused SFPQ degradation through ubiquitin-proteasomal pathway. (A) EA.hy926 cells exposed to TGF-β followed by co-immunoprecipitation with SFPQ and immunoblotted with ubiquitin antibody. (B) EA.hy926 cells were treated with MG132 (0.5uM), and in combination with TGF-β at early time intervals (12hr). Immunoblot analysis of EA.hy926 cell lysates, were probed for SFPQ and PAN-ubiquitin antibodies, and along with their respective densitometry quantification analysis. All analyzed data were normalized to the control treatment condition. (C) EA.hy926 cells were treated with BafilomycinA1 (70uM), and in combination with TGF-β at early time intervals (12hr). Immunoblot analysis of EA.hy926 cell lysates, were probed for SFPQ, and along with their respective densitometry quantification analysis. All analyzed data were normalized to the control treatment condition. (D) EA.hy926 cells were treated with TGF-β for 1 hour and immunoprecipitated for endogenous SFPQ, fresh lysates were probed for K48 ubiquitin linkage along with the immunoblots of the input samples. Values represent the mean ± SD. * p < 0.05, ** p < 0.01, *** p < 0.001 and # p < 0.0001 by unpaired t test or one way ANOVA.

### Nuclear localization of SFPQ decreased upon TGF-β treatment by promoting cytosolic localization without altering its tyrosine phosphorylation

In neuronal cells, cytosolic version of SFPQ abolishes motor axonal defects, rescuing key transcripts, and restores motility in the paralyzed SFPQ null mutants, indicating a non-nuclear processing role in motor axons [24]. Previously, localization of SFPQ were reported to be governed by its post-translational tyrosine phosphorylation [25]. We assessed the total tyrosine phosphorylation status of SFPQ upon stimulation with TGF-β for 1 h. Total tyrosine phosphorylation of SFPQ protein was modestly diminished upon TGF-β treatment, however, such reduction remained statistically insignificant (Figure S2B). Therefore, we next assessed the localization of SFPQ protein in EC exposed to TGF-β. Immunofluorescence staining showed loss of nuclear SFPQ in as early as 12 h post-TGF-β stimulation which persisted at least until 24 h post TGF-β challenge (Figure 4A). Interestingly, at 12 h post-TGF-β exposure, we observed a loss of SFPQ abundance in the core part of the nucleus with a relatively high abundance of remaining SFPQ present close to the nuclear membrane, indicating site-specific reduction of SFPQ protein with TGF-β *in vitro* (Figure 4A). However, we failed to detect the cytosolic localization of SFPQ in any of these treatment groups. We then also performed confocal imaging using EC exposed to TGF-β for 1 h. Interestingly, we could detect compartmentalization of SFPQ localization within the nucleus of EC exposed to TGF-β (Figure S3A), however, in the similar exposure (where we observed the compartmentalization of nuclear SFPQ), we were unable to detect presence of SFPQ in cytosol. We, then, enhanced the exposure which revealed the cytosolic localization of a portion of cellular SFPQ in cells exposed to TGF-β which could not be detected in control cells (Figure S3B).

**Figure 4.**
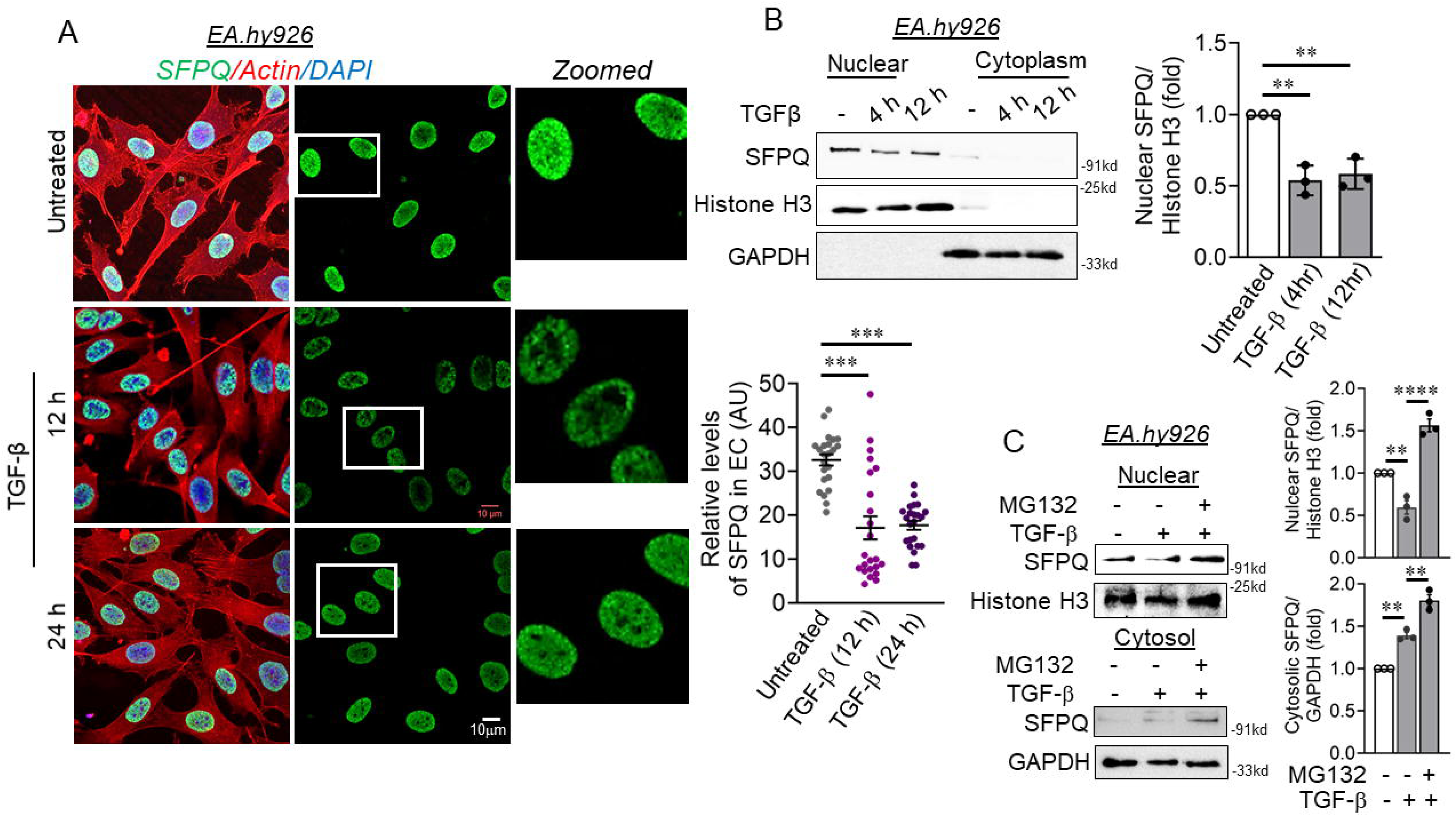
TGF-β caused loss of nuclear localization of SFPQ protein. (A) Immunofluorescence staining of EA.hy926 cells exposed to TGF-β for SFPQ (Green) and Actin filaments (Phalloidin, Red). DAPI staining to visualize the nucleus is shown in blue. Nuclear SFPQ fluorescence signal in individual EA.hy926 cells (dots) from three individual experiments. Fluorescence intensity AU values per individual cells are indicated together with the mean. Total number of cells, n ≥ 30. (B) EA.hy926 cells were treated with TGF-β and separated for cellular components, nuclear (10ug) and cytoplasmic fraction (30ug) were loaded and immunoblot for SFPQ protein, at time intervals of 4, and 12 h. GAPDH and histone H3 showed the purity of cytosolic and nuclear fractions respectively. (C) Cell Fractionation followed by immunoblot analysis was performed using EA.hy926 cells pre-treated with MG132 followed by stimulating with TGF-β for 4 h. Nuclear (10ug) and cytoplasmic fraction (30ug) were loaded and immunoblotted for SFPQ protein. Values represent the mean ± SD. ** p < 0.01, *** p < 0.001 and # p < 0.0001 by unpaired t test.

To confirm the immunofluorescence data, we next investigated the localization of SFPQ using sub-cellular fractionation coupled with immunoblot analysis. We immunoblotted nuclear and cytoplasmic fraction of cell lysates exposed to TGF-β and observed reduction in nuclear SFPQ in cells treated with TGF-β for 4 and 12 h (Figure 4B). We further performed time-dependent cytosolic localization of SFPQ in endothelial cells upon TGF-β treatment for 0.5 and 1 h. Such experiment showed cytosolic localization of a small proportion of SFPQ within 0.5 h of TGF-β treatment which further enhanced upon 1 h of TGF-β challenge (Figure S3C). Moreover, we also detected loss of nuclear SFPQ in EC exposed to TGF-β for 72 h (Figure S3D). Interestingly, similar to immunofluorescence analysis, we were unable to detect the presence of SFPQ in cytoplasmic fraction of EC exposed to these different time points. Therefore, we next assessed the localization of SFPQ in nuclear and cytosolic fractions in EC pretreated with proteasomal inhibitor of MG-132 followed by TGF-β exposure for 4 h. As observed previously, TGF-β caused a reduction in nuclear localization of SFPQ, while MG-132 modestly enhanced nuclear SFPQ level (Figure 4C). Interestingly, we could detect the presence of SFPQ in cytosolic fraction of EC co-treated with MG132 and TGF-β (Figure 4C), thus indicating that TGF-β caused cytosolic translocation of SFPQ prior to its degradation through ubiquitination-dependent proteasomal degradation. These results suggest that SFPQ protein abundance in the nucleus is lost in EC upon TGF-β challenge and further hinting towards a possible role of SFPQ in regulating nuclear associated signaling pathways. Therefore, both of our cell fractionation-immunoblot experiment and immunofluorescence-confocal imaging data suggested that a modest fraction of SFPQ translocate to the cytosol upon TGF-β exposure.

We next questioned whether cytosolic localization of SFPQ upon TGF-β induction was dependent on Smad activation. We therefore assessed the localization of SFPQ in EC which are pre-exposed to pharmacological inhibitor of Smad phosphorylation and induced with TGF-β for 1 h. Through such assessment, we observed that inhibition of Smad phosphorylation failed to block the SFPQ cytosolic localization (Figure S3).

### SFPQ level inversely correlates with mesenchymal-specific transcription factor Snail expression *in vitro*

Similar to epithelial-to-mesenchymal transition, EndMT is mediated by core set of transcription factors-Snail (SNAI1), SLUG (SNAI2), TWIST, and ZEB1/2 [2,26]. To understand the potential function of SFPQ in EndMT, we inhibited protein expression of SFPQ in EA.hy926 cells. siRNA-mediated silencing of SFPQ showed nearly 70% reduction in SFPQ protein expression (Figure 5A). Endothelial cells silenced for SFPQ gene expression showed enhanced Snail protein levels (Figure 5A) with exposure to TGF-β. Surprisingly, SFPQ siRNA transfected EC without exposure to TGF-β also showed a significant increase in the levels of Snail protein (Figure 5A). In addition to its influence on Snail, silencing of SFPQ seemingly protected the protein expression of CD144 (VE Cadherin) in EC exposed to TGF-β challenge (Figure S4A). Meanwhile KLF4 is a known regulator of CD144 [27], upon further examination we found negligible effect of SFPQ knock down on KLF4 protein levels (Figure S4A). Unlike CD144, SFPQ knock down in EC exposed to TGF-β did not show any alterations in the expression of other endothelial specific genes such as eNOS (Figure S4B) and CD31 (Figure S4C).

**Figure 5.**
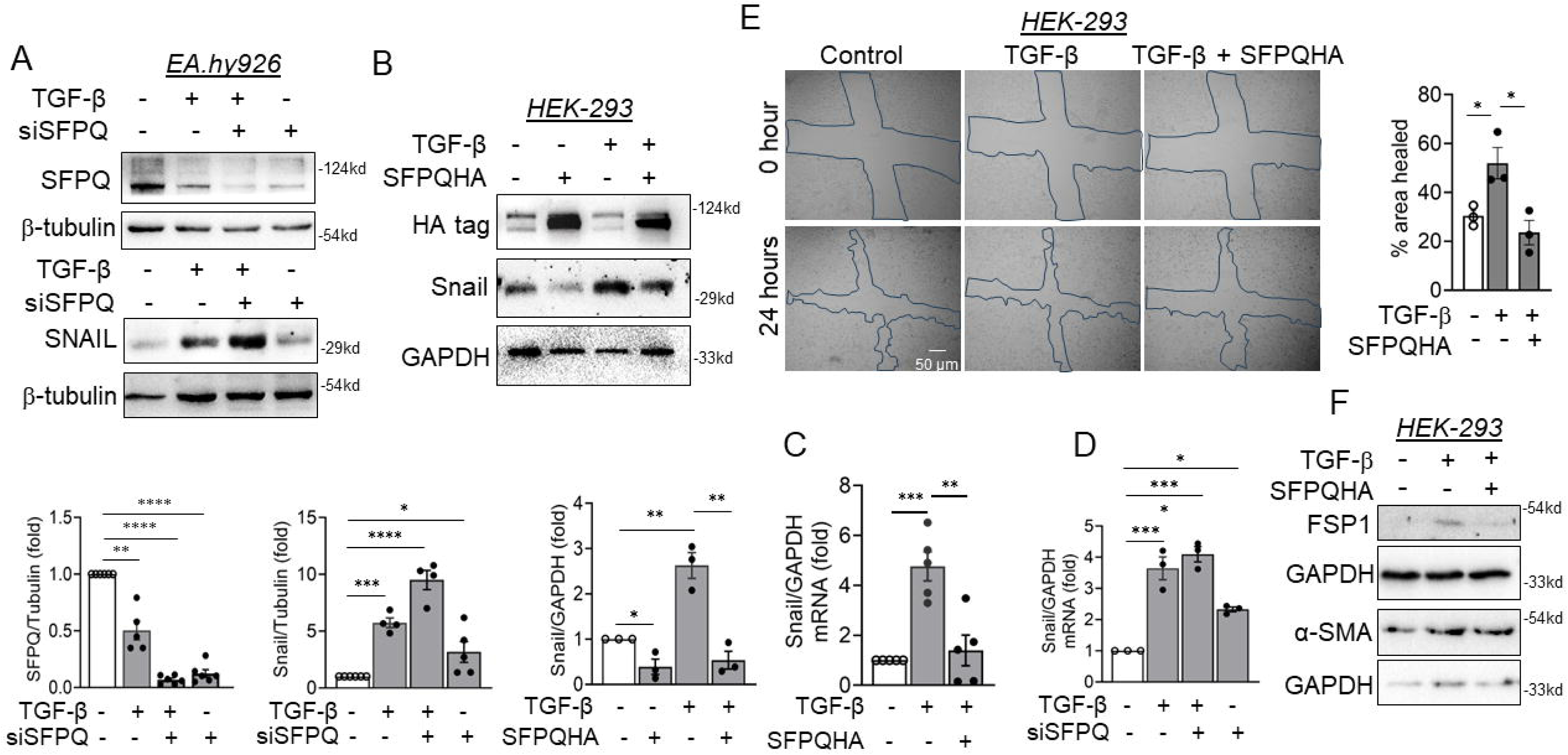
SFPQ protein levels affect the expression of mesenchyme-specific Snail protein. (A) EA.hy926 cells were transfected with SFPQ siRNA, and treated with TGF-β (10 ng/ml) for 72 h, followed by immunoblotting to detect SFPQ and Snail (n = 5). (B) Immunoblot analysis of HEK293 lysates collected from cells transfected with SFPQ overexpression plasmid characterized by expression of HA tag, followed by challenging with TGF-β (10 ng/ml) for 72 h and probed for Snail (n = 4) along with densitometry quantification of the blots. (C) Transcript-level expression of SNAIL (n = 5) measured through RT-qPCR technique in HEK293 expressing SFPQ exogenously and treated with TGF-β condition. All analyzed data were normalized to the control treatment condition. (D) Transcript-level expression of SNAIL (n = 3) measured through RT-qPCR technique in EA.hy926 cells transfected with SFPQ siRNA and treated with TGF-β condition. (E) Wound healing assay using HEK293 cells treated with TGF-β for 24 hours along with HA-SFPQ overexpression. All analyzed data were normalized to the control treatment condition. The microscopy images were captured at 10x magnification and the analysis was performed using ImageJ software. (F) Immunoblots HEK293 cells treated with TGF-β or TGF in combination with SFPQ overexpression. Values represent the mean ± SD. * p < 0.05, ** p < 0.01, *** p < 0.001, and **** p < 0.0001 by unpaired t test.

We next questioned whether an overexpression of SFPQ could reverse TGF-β induced Snail expression. To do so, we used an HA-tagged SFPQ plasmid and transfected the same into HEK293 cells followed by induction with TGF-β. In so doing, we observed that plasmid-mediated exogenous expression of SFPQ in HEK293 cells abrogated TGF-β-induced protein level expression of Snail (Figure 5B). More importantly, forced expression of SFPQ in TGF-β nontreated HEK293 cells also significantly reduced the basal level of Snail expression (Figure 5B). Furthermore, Overexpression of SFPQ in HEK293 cells resulted in the reduction of Snail transcript levels as compared to TGF-β-stimulated non-transfected cells (Figure 5C). We then assessed the transcript level of SLUG, another transcription factor regulated by TGF-β, in HEK293 cells overexpressing SFPQ gene and further exposed to TGF-β. Herein, we observed significant increase in the transcript level of SLUG gene; however, overexpression of SFPQ could not reverse TGF-β-induced SLUG transcript expression (Figure S4D). Such data indicate specific regulation of Snail but not Slug by SFPQ. In total, we demonstrated that SFPQ level counter-regulates Snail gene expression both in TGF-β induced and non-induced experimental conditions. We next performed a wound-healing assay in HEK293 cells subjected to TGF-β treatment alone or in combination with SFPQ overexpression. TGF-β treatment promoted a significant increase in wound closure compared to control cells; however, this effect was attenuated upon SFPQ overexpression, resulting in recovery rates comparable to those of untreated controls, underscoring the role of SFPQ in restraining Snail-driven cell migration under TGF-β conditions (Figure 5E). Consistently, immunoblot analysis of cell lysates revealed elevated levels of FSP1 and α-SMA upon TGF-β stimulation, which were markedly reduced in cells co-treated with TGF-β and SFPQ overexpression (Figure 5F).

### SFPQ is associated with transcription factors regulating Snail gene expression, however, such association remained unaltered upon TGF-β exposure

A very recent study described sequestration of Smad4 by SFPQ condensates which compromised the tumor supportive signaling imparted by TGF-β [28]. Therefore, we questioned whether in the current experimental settings, SFPQ protein associates with different transcriptional regulators such as Smad2/3, Smad4, and N1-ICD, all of which play important role in TGF-β dependent transcriptional expression of Snail. Moreover, we also analyzed whether the association of these transcription factors with SFPQ diminished upon TGF-β induction especially at shorter time point when SFPQ total and nuclear protein remained unchanged. We therefore, performed co-immunoprecipitation experiment using the nuclear fractions to analyze the association of SFPQ with these transcription factor in the nucleus of the cells stimulated with TGF-β for a shorter time point of 1 h. In doing so, we detected robust association of endogenously expressing N1-ICD (Figure 6A), and Smad4 (Figure 6B) in untreated EC. Surprisingly, such association of SFPQ with Smad4 (Figure 6B), and N1-ICD (Figure 6A) remained unchanged in EC stimulated with TGF-β. Moreover, we next performed co-immunoprecipitation analysis using lysates from cells which are treated with TGF-β for 12 h at which time point we observed reduction in SFPQ level. We choose this time point as SFPQ level was observed to be significantly reduced by 12 h post TGF-β exposure. In longer time point, we detected loss of SFPQ association with Smad2/3 suggesting that loss of SFPQ finally caused loss of their association with Smad2/3 (Figure 6C). We also analyzed the association of SFPQ with other nuclear proteins including its conventional interacting partner of the paraspeckle, p54nrb in EC treated with TGF-β. We could not detect any alteration in the association of SFPQ with its known binding partner p54nrb upon TGF-β stimulation (Figure S5A). Importantly, we did not detect any association of SFPQ with other nuclear proteins such as LSD1, and HDAC1 after 1 hour of TGF-β induction (Figure S5A).

**Figure 6.**
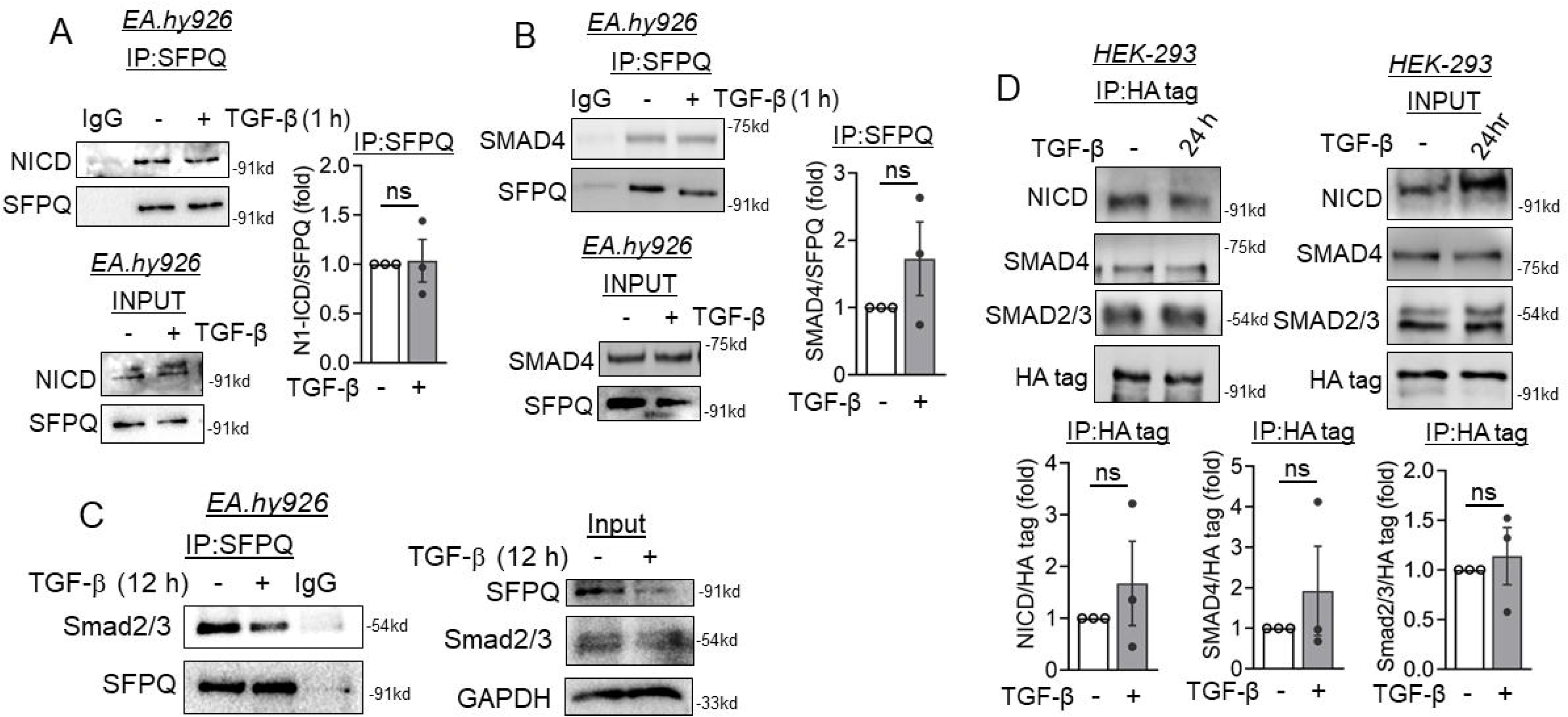
SFPQ interacted with many partners of the TGF-β signaling pathway, but TGF-β stimulation failed to alter such association. (A-B) EA.hy926 cells were treated with TGF-β for 1 h and immunoprecipitated for endogenous SFPQ, fresh lysates were probed for Cleaved Notch1 (NICD) (A) and Smad4 (B) followed by comparing them with Input samples. (C) EA.hy926 cells were treated with TGF-β for 12 h and immunoprecipitated with SFPQ antibody, immunoblotted for Cleaved Smad2/3 followed by comparing them with Input samples. (D) HEK293 cells were transfected with SFPQ overexpression plasmid and treated with TGF-β for 24 h, lysate was subjected to anti-HA tag pulldown followed by immunoblotting for Cleaved Notch1 (NICD), Smad2/3, and Smad4. Values represent the mean ± SD.

We next assessed the association of SFPQ with Smad2/3, Smad4, and N1-ICD in HEK293 cells where HA-tagged SFPQ was forced expressed using plasmid. Parallel to the observation in EC, SFPQ robustly associated with Smad2/3, Smad4, and N1-ICD in untreated cells which remained unaltered even after TGF-β exposure (Figure 6D). These data strongly hint that TGF-β does not alter the association of SFPQ with proteins that are responsible for TGF-β-induced Snail expression. Therefore, unlike the previous study [28], regulation of TGF-β downstream signaling especially its control of Snail gene expression through sequestration of these transcription factors by SFPQ does not hold true in the current experimental settings.

### SFPQ occupied two distinct regions of Snail gene promoter and coding region which likely causes the transcriptional repression of Snail

Because we could not detect any alteration in the association of Smad2/3, Smad4, and N1-ICD with SFPQ upon TGF-β challenge, we therefore assessed the possibility of SFPQ being a transcriptional repressor of Snail gene expression by binding to its gene promoter/coding region. Numerous reports have demonstrated SFPQ to act as a transcriptional repressor/activator by binding directly to the chromatin [8,29]. To comprehensively decipher the occupancy of SFPQ on Snail promoter region, we performed Chromatin immunoprecipitation assay followed by qPCR analysis. We designed multiple primers spanning Transcription Start Site (TSS) and upstream/downstream to TSS (−700bp to +800bp) (Figure 7A). Interestingly, we detected a pronounced binding of SFPQ to region ∼700bp downstream to the TSS, in untreated EC (Figure 7B). We also detected a modest binding of SFPQ in the proximal promoter region of Snail gene (Figure 7B, ∼645 bp upstream to TSS) near the E-box promoter region where many transcriptional complexes including Smad proteins binds to initiate *SNAIL* gene transcription.

**Figure 7.**
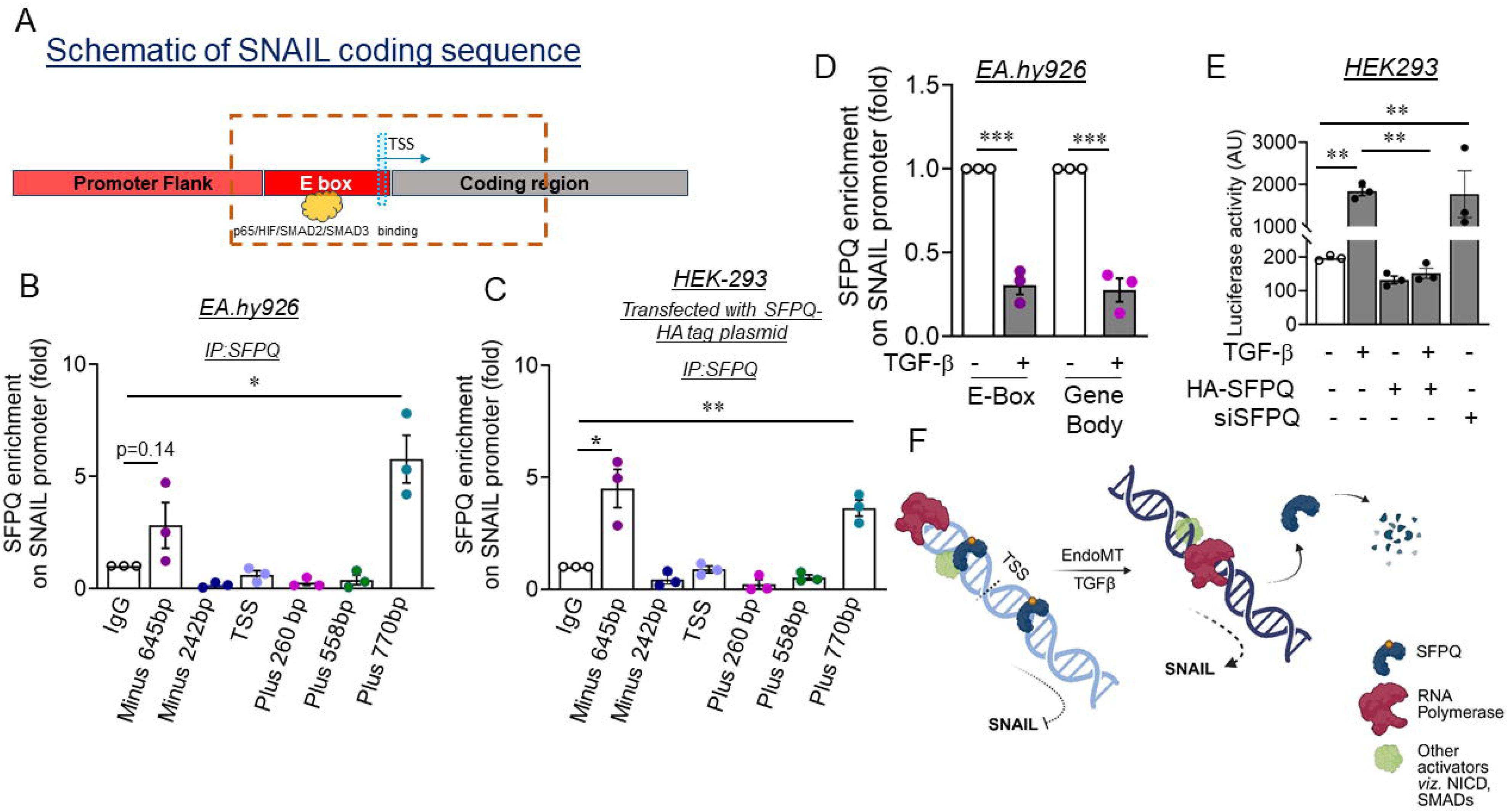
SFPQ occupancy study showed preferential binding at SNAIL coding region 700bp downstream to TSS. (A) Schematic diagram of SNAIL promoter sequence and potential binding sites for transcriptional activators viz. NFκB, HIF, SMADs. Chromatin occupancy study by Cut-and-run assay in EA.hy926 cell lysates and HEK293 lysates overexpressing SFPQ were pulldown with endogenously (B) and exogenously (C) expressed SFPQ, followed by qPCR of eluent DNA with multiple primers spanning the SNIAL promoter region. All Ct values were normalized to the Ct value of control IgG. (D) Chromatin occupancy study by Cut-and-run assay in EA.hy926 cell treated with TGF-β for 4 h using SFPQ followed by qPCR of eluent DNA with primers for the E-Box region and gene coding region of SNAIL gene. All Ct values were normalized to the Ct value of control IgG. (F) In endothelial cells, SFPQ is expressed constitutively and is associated with multiple nuclear processes. SFPQ binds both upstream and downstream to TSS of Snail gene, thereby restricting the transcriptional activity of Snail transcription factors such as Smad2/3, Smad4, and N1-ICD by associating with them and also essentially averting transcriptional activation of SNAIL. With cytokine TGF-β stimulus, endothelial cells reduce abundance of nuclear SFPQ by degrading it post translationally. This reduction in SFPQ levels in the nucleus results in the loss of transcriptional repression activity, thereby promoting SNAIL expression.

We next assessed the same binding status of SFPQ protein in *SNAIL* promoter/coding region in SFPQ overexpressing HEK293 cells. In SFPQ overexpressing HEK-293 cells, we observed a robust binding of SFPQ to the same region (Figure 7C, ∼700bp downstream to TSS). Moreover, in these experimental settings, we detected robust binding of SFPQ in the proximal promoter region (E-box region) of SNAIL gene (Figure 7C, ∼645 bp upstream to TSS). These results suggest potential binding of SFPQ to SNAIL-encoding DNA to govern binding of other activating factors, averting activation of Snail gene in both EC and HEK293 cells. In addition, we also performed the CUT&RUN assay using endothelial cells treated with TGF-β for 4 h in which time point we observed significant loss of SFPQ protein level. Through such experiment, we observed significant loss of SFPQ enrichment in both E-Box and gene body region of the Snail gene (Figure 7D). We also performed the ChIP assay using Smad2/3 antibody in cells where SFPQ was knockdown. Interestingly, loss of SFPQ in TGF-β treated endothelial cells showed modest increase in Smad2/3 enrichment on the E-box region of the Snail promoter (Figure S5B). This indicate that even though, SFPQ protein associates with Smad2/3, however, Smad2/3 capacity to bind to Snail promoter remained unaffected by SFPQ knockdown.

We next performed reporter assay using Snail_pGL2 plasmid containing the Snail promoter (−1045 to −56, which harbors the E-box region of the promoter, which we showed to be bound by SFPQ for the transcriptional repression) sequence upstream of luciferase. Using the same, we confirmed that HEK293 cells treated with TGF-β exhibited nearly 9-fold enhanced luciferase activity compared to control while overexpression of HA tagged SFPQ caused complete reversal of TGF-β induced luciferase activity (Figure 7E). Moreover, knockdown of SFPQ in cells caused significant (equivalent to TGF-β) increase in luciferase activity (Figure 7E). Altogether, these data suggest that SFPQ acts as a transcriptional repressor by binding to the E-box region. However, the employed construct does not contain the coding region which we showed to be bound by SFPQ and therefore the significance of SFPQ binding to this region and Snail transcriptional repression is yet to be validated.

## Discussion

TGF-β, a pleiotropic cytokine, is known to be differentially involved in the regulation of multi-lineage differentiation of stem cells, through the Smad and non-Smad pathways [30]. Here, we identified that SFPQ, a DNA- and RNA-binding protein acts as a transcriptional repressor of Snail, a TGF-β signaling-regulated transcription factor, through enrichment at proximal promoter/coding region of SNAIL gene. We reported loss of endothelial SFPQ in kidney tissues of rats that underwent subtotal nephrectomy or cultured EC treated with TGF-β. Moreover, loss of SFPQ was associated with an increase in Snail expression. Upon TGF-β stimulation, SFPQ lost its nuclear localization, translocated to the cytosol, followed by degradation via ubiquitin-proteasomal pathway. Using knock down and forced expression approach, we next described that SFPQ level oppositely correlate with the level of Snail in different cell types, both in absence and presence of TGF-β. Alteration in SFPQ did not affect the expression of other TGF-β associated transcription factor, Slug. Interestingly, we found that SFPQ associated with many TGF-β regulatory transcription factors including Smad2/3, Smad4, and N1-ICD, however, such association remained unchanged upon TGF-β induction. Finally, we showed that SFPQ enriched in the proximal promoter region (E-box region) and initial coding region of SNAIL gene which likely caused transcriptional repression of SNAIL gene expression (Figure 7F).

Emerging evidence shows that the phenomenon of EndMT is regulated by several non-primary contributors, such as non-coding RNAs, accessory transcription factors, epigenetic modifications, and RNA binding proteins [31,32]. A study has established that ectopic expression of miR-20a inhibits TGF-β1-mediated EndMT induction. Furthermore, they demonstrated FGF2 inadvertently increases expression of miR-20a in endothelial cells and abrogates endothelial responsiveness to TGF-β1 [15]. RASAL1 expression has been well studied in progression of fibrosis involving kidney and liver [33], DNMT1 and HDAC2 acts in synergy to maintain hypermethylated repressive state of RASAL1 promoter, ensuing enhanced Ras-GTP activity and in the activation of transcription factors Snail, Twist, and Slug [34]. Previous study from our lab reported reported that enhanced H3K4me3 caused transcriptional activation of Notch-associated ligands-Jagged1/2. This aberrant expression of Notch ligands further triggered activation of Notch receptor intracellular domain (NICD) in the nucleus and encoded EndMT specific factors-Slug, and alpha smooth muscle actin, imparting mesenchymal signature to ECs [35]. New studies are advancing to establish the significance of RNA binding proteins (RBP) in maintaining cellular homeostasis. Liang et. al., proved that RBP LARP7 interacts with epigenetic repressor TRIM28 (tripartite motif containing 28) and colocalizes at the SLUG promoter. This complex in natural state repressed transcription of SLUG through deacetylating the histones encoding it. Moreover, inducible knockout of LARP7 and/or TRIM28 in the endocardium accelerated EndMT subsequently resulting in the valvular hyperplasia [36]. SFPQ is one of the ubiquitously expressed RNA-binding proteins involved in RNA transport, apoptosis, DNA damage repair, genome stability, and innate immune response [37]. Here in this study, we showed that in normal EC, the SFPQ binds to the SNAIL coding and proximal promoter region and represses SNAIL transcription. However, when EC are exposed to pro-EndMT signals like TGF-β, they downregulate SFPQ thereby impairing the recruitment of SFPQ to chromatin, leading to the loss of SNAIL repression. Interestingly, inhibition of TGF-β signaling through pharmacological inhibitors of Smads phosphorylation by SIS3 reversed SFPQ protein level. While SIS3-mediated inhibition of Smad3 suppressed Snail induction, interestingly, it did not affect the cytosolic translocation of SFPQ, indicating that SFPQ relocalization occurs independently of canonical Smad3 signaling. Furthermore, the ability of TGF-β to further enhance Snail expression even in SFPQ knockdown conditions supports the involvement of an additional, SFPQ-independent mechanism. These data strongly hint at a dual regulatory role of TGF-β in Snail induction, highlighting both SFPQ-dependent and independent pathways as part of its broader transcriptional regulatory network.

Tyrosine phosphorylation of SFPQ has also been linked to aberrant cytoplasmic localization in cancer cells. In anaplastic large-cell lymphomas (ALCLs) expressing the NPM/ALK fusion protein, SFPQ is phosphorylated on tyrosine 293 within the PR linker by the ALK kinase domain. This tyrosine phosphorylation causes the export of SFPQ from the nucleus to the cytoplasm [25]. Similarly, the tyrosine kinase BRK (breast tumor kinase) phosphorylates SFPQ in response to EGF signaling in breast cancer cells, leading to cytoplasmic localization of SFPQ. The specific BRK phosphorylation site has not been determined, but it likely occurs within or near the SFPQ nuclear localization signal (NLS) in the C-terminus [38]. These findings suggest tyrosine phosphorylation may represent a regulatory mechanism to control SFPQ localization and activity, not only in cancer but also potentially in normal cells as well. However, limited studies have explored the possibility of dephosphorylation of SFPQ protein. Studies have demonstrated an important role for the phosphatase protein phosphatase 1 (PP1) in regulating SFPQ. The C-terminus of SFPQ’s first RNA recognition motif (RRM1) contains an RVxF sequence, which is the consensus binding motif for PP1 [39]. Consistent with this, SFPQ has been shown to interact with PP1 through co-immunoprecipitation and co-localization studies, though this interaction appears to be indirect and requires the DBHS protein p54nrb/NONO. PP1 can directly dephosphorylate SFPQ both *in vitro* in cell free system and in cells. Inhibition of PP1 by ceramide leads to increased phosphorylation of SFPQ, indicating PP1 activity normally counteracts the phosphorylation of SFPQ [40]. Nonetheless, these findings demonstrate the importance of balanced phosphorylation and dephosphorylation in controlling the diverse functions of the multifunctional protein SFPQ. We discovered that the loss of nuclear SFPQ is an early event; immunofluorescence imaging showed TGF-β stimulus depletes SFPQ abundance in a site-specific manner. Interestingly, we confirmed that such changes in SFPQ localization upon TGF-β induction was independent of its post translational modification through tyrosine phosphorylation. One of the possible explanations to this is that SFPQ is involved as a part of spliceosome complex that works closely with RNA Polymerase II [41], being a part active transcriptional machinery, and eventual degradation of SFPQ (TGF-β exposure) results in global loss of SFPQ clusters. In addition, the application of zinc to primary cortical neurons induced the cytoplasmic accumulation and aggregation of SFPQ. Mutagenesis of the three zinc-coordinating histidine residues resulted in a significant reduction in the zinc-binding affinity of SFPQ in solution and the zinc-induced cytoplasmic aggregation of SFPQ in cultured neurons. Therefore, zinc availability likely represents a mechanism for the imbalance in the nucleocytoplasmic distribution of SFPQ [42]. Interestingly, TGF-β signaling can also regulate the expression and activity of zinc transporters and metallothioneins, which are proteins involved in zinc homeostasis [43]. Moreover, TGF-β stimulation can lead to an increase in intracellular zinc levels, particularly in CD4+ T cells [44]. Therefore, in the current study, TGF-β mediated regulation of intracellular zinc could influence the cytosolic localization of SFPQ protein, however, such hypothesis need to be tested experimentally in current experimental settings.

Notch1 has notably been identified as a mechanosensor playing a crucial role in promoting and maintaining arterial homeostasis [45]. Furthermore, non-canonical Notch complex regulates adherens junctions and maintains endothelial barrier integrity in microvessels under flow conditions [46]. TGF-β-mediated SMAD3 complex is reported to activate HEY1 gene and contribute to Epithelial-to-mesenchymal transition (EMT). This indicates TGF-β/ NOTCH crosstalk coordinate activated state of downstream genes driving cells to EMT [47]. The widespread expression of NICD in endothelial and endocardial cells is also reported to trigger the expression of pro-EMT and mesenchymal genes, including SNAI1, SNAI2, TGF-β2, TWIST2, and Alk3 [48]. In our study, in unstimulated conditions, nuclear SFPQ co-immunoprecipitated with Cleaved Notch 1 protein (N1-ICD), Smad2/3, and Smad4. However, short-term or long-term exposure to TGF-β in EC and HEK293 cells, respectively, failed to modify the association of these transcription factors with SFPQ.

SFPQ knockdown in endothelial cells displayed enhanced expression of Snail protein levels, and the effect was exaggerated in combination with TGF-β. This phenotype strongly suggests the contribution of SFPQ in regulating SNAIL expression. Remarkably, upon expressing SFPQ exogenously we observed a robust reversal of SNAIL protein/transcript expression in TGF-β challenged HEK-293 cells. This evidence confirms that SFPQ is an upstream regulator of the transcription factor SNAIL. We also demonstrated that SFPQ knockdown in EC enhanced expression of endothelial specific protein VE-cadherin (CD144).Rudini et al., showed inactivation or absence of VE-cadherin blunts the responsiveness of ECs to TGF-β, and co-immunoprecipitation experiments confirmed VE-cadherin interacts with TβRII, ALK5 and ALK1 and assembles into an active receptor complex [49]. The hypothesis that loss of SFPQ may enhance the expression of VE-cadherin to heighten the effect of TGF-β exposure in endothelial cells requires further investigation. SFPQ occupancy studies on SNAIL regulatory regions suggested a weak binding of SFPQ to SNAIL proximal promoter region (∼645 bp upstream to TSS) while robust binding in coding region (∼700bp downstream to TSS). One hypothesis could be that SFPQ may act as a repressor to stall activity of RNA Polymerase; thus, loss of SFPQ enhanced the expression of SNAIL, and following exogeneous overexpression of SFPQ resulted in robust reduction in SNAIL transcripts. However, more experiments are required to provide evidence that SFPQ act as a transcriptional repressor of SNAIL gene expression.

In conclusion, this study highlights the contribution of RNA binding protein SFPQ in regulating expression of SNAIL protein. Absence of SFPQ in response to pro-inflammatory cytokine TGF-β results in the reduction in nuclear SFPQ abundance and a loss of enrichment on SNAIL proximal promoter and coding region. Post translational modification through ubiquitination of SFPQ protein propels SFPQ towards degradation causing loss of SFPQ binding to the SNAIL promoter/coding region. Future studies exploring the role of SFPQ in organ fibrosis induced by TGF-β signaling will allow further understanding of how these RNA binding proteins like SFPQ governs onset and progression of human disease.

## Acknowledgment

This work was supported by the Core Research Grant from Anusandhan National Research Foundation (formerly Science and Engineering Research Board)-Department of Science and Technology, Govt. of India (CRG/2022/002209) to SM. This work was also partly supported by a Competitive Research Grant from the Department of Biotechnology, Govt. of India (BT/PR33144/MED/30/2170/2019) to SM. N.P.T. was supported by a graduate fellowship from BITS Pilani. S.R. is supported by a graduate fellowship from Department of Science and Technology-Innovation in Science Pursuit for Inspired Research fellowship (DST/INSPIRE/03/2019/000582). Y.T.K. is supported by a graduate fellowship from BITS Pilani. A.A. holds the Keenan Chair in Medicine from St. Michael’s Hospital and University of Toronto. The histological studies were supported, in part, by a project grant from the Canadian Institutes of Health Research to A.A. (PJT166083) and through a John R. Evans Leaders Fund Award from the Canada Foundation for Innovation (38214). We gratefully acknowledge the technical assistance of Mr. Suman Kumar (Confocal Facility, BITS Pilani, Pilani Campus).

## Duality of Interest Statement

A.A. has received research support through his institution from Boehringer Ingelheim.

## References

[1] Y. Yoshimatsu, T. Watabe, Emerging roles of inflammation-mediated endothelial– mesenchymal transition in health and disease, Inflamm. Regen. 42 (2022) 9. 10.1186/s41232-021-00186-3.

[2] J.L. Wrana, L. Attisano, R. Wieser, F. Ventura, J. Massagué, Mechanism of activation of the TGF-β receptor, Nature 370 (1994) 341–347. 10.1038/370341a0.

[3] H. Fujita, M. Kang, M. Eren, L.A. Gleaves, D.E. Vaughan, T. Kume, Foxc2 Is a Common Mediator of Insulin and Transforming Growth Factor β Signaling to Regulate Plasminogen Activator Inhibitor Type I Gene Expression, Circ. Res. 98 (2006) 626–634. 10.1161/01.RES.0000207407.51752.3c.

[4] T. Vincent, E.P.A. Neve, J.R. Johnson, A. Kukalev, F. Rojo, J. Albanell, K. Pietras, I. Virtanen, L. Philipson, P.L. Leopold, R.G. Crystal, A.G. De Herreros, A. Moustakas, R.F. Pettersson, J. Fuxe, A SNAIL1–SMAD3/4 transcriptional repressor complex promotes TGF-β mediated epithelial–mesenchymal transition, Nat. Cell Biol. 11 (2009) 943–950. 10.1038/ncb1905.

[5] H.A.R. Hadi, J.A. Suwaidi, Endothelial dysfunction in diabetes mellitus, Vasc. Health Risk Manag. 3 (2007) 853–876.

[6] W.H. Hudson, E.A. Ortlund, The structure, function and evolution of proteins that bind DNA and RNA, Nat. Rev. Mol. Cell Biol. 15 (2014) 749–760. 10.1038/nrm3884.

[7] T. Hirose, G. Virnicchi, A. Tanigawa, T. Naganuma, R. Li, H. Kimura, T. Yokoi, S. Nakagawa, M. Bénard, A.H. Fox, G. Pierron, NEAT1 long noncoding RNA regulates transcription via protein sequestration within subnuclear bodies, Mol. Biol. Cell 25 (2014) 169–183. 10.1091/mbc.e13-09-0558.

[8] K. Imamura, N. Imamachi, G. Akizuki, M. Kumakura, A. Kawaguchi, K. Nagata, A. Kato, Y. Kawaguchi, H. Sato, M. Yoneda, C. Kai, T. Yada, Y. Suzuki, T. Yamada, T. Ozawa, K. Kaneki, T. Inoue, M. Kobayashi, T. Kodama, Y. Wada, K. Sekimizu, N. Akimitsu, Long Noncoding RNA NEAT1-Dependent SFPQ Relocation from Promoter Region to Paraspeckle Mediates IL8 Expression upon Immune Stimuli, Mol. Cell 53 (2014) 393–406. 10.1016/j.molcel.2014.01.009.

[9] S.-W. He, C. Xu, Y.-Q. Li, Y.-Q. Li, Y. Zhao, P.-P. Zhang, Y. Lei, Y.-L. Liang, J.-Y. Li, Q. Li, Y. Chen, S.-Y. Huang, J. Ma, N. Liu, Correction: AR-induced long non-coding RNA LINC01503 facilitates proliferation and metastasis via the SFPQ-FOSL1 axis in nasopharyngeal carcinoma, Oncogene 40 (2021) 6703–6704. 10.1038/s41388-021-02050-7.

[10] H. Qin, H. Ni, Y. Liu, Y. Yuan, T. Xi, X. Li, L. Zheng, RNA-binding proteins in tumor progression, J. Hematol. Oncol.J Hematol Oncol 13 (2020) 90. 10.1186/s13045-020-00927-w.

[11] Y. Fukuda, M.F. Pazyra-Murphy, E.S. Silagi, O.E. Tasdemir-Yilmaz, Y. Li, L. Rose, Z.C. Yeoh, N.E. Vangos, E.A. Geffken, H.-S. Seo, G. Adelmant, G.H. Bird, L.D. Walensky, J.A. Marto, S. Dhe-Paganon, R.A. Segal, Binding and transport of SFPQ-RNA granules by KIF5A/KLC1 motors promotes axon survival, J. Cell Biol. 220 (2021) e202005051. 10.1083/jcb.202005051.

[12] N. Fujita, D.L. Jaye, M. Kajita, C. Geigerman, C.S. Moreno, P.A. Wade, MTA3, a Mi-2/NuRD complex subunit, regulates an invasive growth pathway in breast cancer, Cell 113 (2003) 207–219. 10.1016/s0092-8674(03)00234-4.

[13] A. Advani, K.A. Connelly, D.A. Yuen, Y. Zhang, S.L. Advani, J. Trogadis, M.G. Kabir, E. Shachar, M.A. Kuliszewski, H. Leong-Poi, D.J. Stewart, R.E. Gilbert, Fluorescent Microangiography Is a Novel and Widely Applicable Technique for Delineating the Renal Microvasculature, PLoS ONE 6 (2011) e24695. 10.1371/journal.pone.0024695.

[14] K. Thieme, S. Majumder, A.S. Brijmohan, S.N. Batchu, B.B. Bowskill, T.A. Alghamdi, S.L. Advani, M.G. Kabir, Y. Liu, A. Advani, EP4 inhibition attenuates the development of diabetic and non-diabetic experimental kidney disease, Sci. Rep. 7 (2017) 3442. 10.1038/s41598-017-03237-3.

[15] A.C.P. Correia, J.-R.A.J. Moonen, M.G.L. Brinker, G. Krenning, FGF2 inhibits endothelial–mesenchymal transition through microRNA-20a-mediated repression of canonical TGF-β signaling, J. Cell Sci. 129 (2016) 569–579. 10.1242/jcs.176248.

[16] G. Kökény, Á. Németh, J.B. Kopp, W. Chen, A.J. Oler, A. Manzéger, L. Rosivall, M.M. Mózes, Susceptibility to kidney fibrosis in mice is associated with early growth response-2 protein and tissue inhibitor of metalloproteinase-1 expression, Kidney Int. 102 (2022) 337–354. 10.1016/j.kint.2022.03.029.

[17] H. Zheng, M. Shen, Y.-L. Zha, W. Li, Y. Wei, M.A. Blanco, G. Ren, T. Zhou, P. Storz, H.-Y. Wang, Y. Kang, PKD1 phosphorylation-dependent degradation of SNAIL by SCF-FBXO11 regulates epithelial-mesenchymal transition and metastasis, Cancer Cell 26 (2014) 358–373. 10.1016/j.ccr.2014.07.022.

[18] V.M. Díaz, A.G. de Herreros, F-box proteins: Keeping the epithelial-to-mesenchymal transition (EMT) in check, Semin. Cancer Biol. 36 (2016) 71–79. 10.1016/j.semcancer.2015.10.003.

[19] K.-S. Hong, K.-J. Ryu, H. Kim, M. Kim, S.-H. Park, T. Kim, J.W. Yang, C. Hwangbo, K.D. Kim, Y.-J. Park, J. Yoo, MSK1 promotes colorectal cancer metastasis by increasing Snail protein stability through USP5-mediated Snail deubiquitination, Exp. Mol. Med. 57 (2025) 820– 835. 10.1038/s12276-025-01433-0.

[20] V.M. Díaz, R. Viñas-Castells, A. García de Herreros, Regulation of the protein stability of EMT transcription factors, Cell Adhes. Migr. 8 (2014) 418–428. 10.4161/19336918.2014.969998.

[21] A. García de Herreros, Control of Snail1 protein stability by post-translational modifications: the basis for a complex regulation of Snail1 function, Int. J. Biol. Sci. 21 (2025) 3183–3196. 10.7150/ijbs.108903.

[22] S. Nakagawa, K. Nishihara, H. Miyata, H. Shinke, E. Tomita, M. Kajiwara, T. Matsubara, N. Iehara, Y. Igarashi, H. Yamada, A. Fukatsu, M. Yanagita, K. Matsubara, S. Masuda, Molecular Markers of Tubulointerstitial Fibrosis and Tubular Cell Damage in Patients with Chronic Kidney Disease, PloS One 10 (2015) e0136994. 10.1371/journal.pone.0136994.

[23] M. Jinnin, H. Ihn, K. Tamaki, Characterization of SIS3, a novel specific inhibitor of Smad3, and its effect on transforming growth factor-beta1-induced extracellular matrix expression, Mol. Pharmacol. 69 (2006) 597–607. 10.1124/mol.105.017483.

[24] S. Thomas-Jinu, P.M. Gordon, T. Fielding, R. Taylor, B.N. Smith, V. Snowden, E. Blanc, C. Vance, S. Topp, C.-H. Wong, H. Bielen, K.L. Williams, E.P. McCann, G.A. Nicholson, A. Pan-Vazquez, A.H. Fox, C.S. Bond, W.S. Talbot, I.P. Blair, C.E. Shaw, C. Houart, Non-nuclear Pool of Splicing Factor SFPQ Regulates Axonal Transcripts Required for Normal Motor Development, Neuron 94 (2017) 931. 10.1016/j.neuron.2017.04.036.

[25] A. Galietta, R.H. Gunby, S. Redaelli, P. Stano, C. Carniti, A. Bachi, P.W. Tucker, C.J. Tartari, C.-J. Huang, E. Colombo, K. Pulford, M. Puttini, R.G. Piazza, H. Ruchatz, A. Villa, A. Donella-Deana, O. Marin, D. Perrotti, C. Gambacorti-Passerini, NPM/ALK binds and phosphorylates the RNA/DNA-binding protein PSF in anaplastic large-cell lymphoma, Blood 110 (2007) 2600–2609. 10.1182/blood-2006-01-028647.

[26] P.-Y. Chen, L. Qin, M. Simons, TGFβ signaling pathways in human health and disease, Front. Mol. Biosci. 10 (2023) 1113061. 10.3389/fmolb.2023.1113061.

[27] C.E. Cowan, E.E. Kohler, T.A. Dugan, M.K. Mirza, A.B. Malik, K.K. Wary, Krüppel-Like Factor-4 Transcriptionally Regulates VE-Cadherin Expression and Endothelial Barrier Function, Circ. Res. 107 (2010) 959–966. 10.1161/CIRCRESAHA.110.219592.

[28] M. Xiao, F. Wang, N. Chen, H. Zhang, J. Cao, Y. Yu, B. Zhao, J. Ji, P. Xu, L. Li, L. Shen, X. Lin, X.-H. Feng, Smad4 sequestered in SFPQ condensates prevents TGF-β tumor-suppressive signaling, Dev. Cell 59 (2024) 48–63.e8. 10.1016/j.devcel.2023.11.020.

[29] E. Rosonina, J.Y.Y. Ip, J.A. Calarco, M.A. Bakowski, A. Emili, S. McCracken, P. Tucker, C.J. Ingles, B.J. Blencowe, Role for PSF in Mediating Transcriptional Activator-Dependent Stimulation of Pre-mRNA Processing In Vivo, Mol. Cell. Biol. 25 (2005) 6734– 6746. 10.1128/MCB.25.15.6734-6746.2005.

[30] M.-K. Wang, Different roles of TGF-β in the multi-lineage differentiation of stem cells, World J. Stem Cells 4 (2012) 28. 10.4252/wjsc.v4.i5.28.

[31] Q. Jun, L. Youhong, Z. Yuan, Y. Xi, B. Wang, S. Xinyi, Y. Fu, C. Kedan, J. Lian, Z. Jianqing, Histone modification of endothelial-mesenchymal transition in cardiovascular diseases, Front. Cardiovasc. Med. 9 (2022) 1022988. 10.3389/fcvm.2022.1022988.

[32] M.S. Hulshoff, G. Del Monte-Nieto, J. Kovacic, G. Krenning, Non-coding RNA in endothelial-to-mesenchymal transition, Cardiovasc. Res. 115 (2019) 1716–1731. 10.1093/cvr/cvz211.

[33] X. Xu, X. Tan, B. Tampe, G. Nyamsuren, X. Liu, L.S. Maier, S. Sossalla, R. Kalluri, M. Zeisberg, G. Hasenfuss, E.M. Zeisberg, Epigenetic balance of aberrant Rasal1 promoter methylation and hydroxymethylation regulates cardiac fibrosis, Cardiovasc. Res. 105 (2015) 279–291. 10.1093/cvr/cvv015.

[34] X. Tan, X. Xu, M. Zeisberg, E.M. Zeisberg, DNMT1 and HDAC2 Cooperate to Facilitate Aberrant Promoter Methylation in Inorganic Phosphate-Induced Endothelial-Mesenchymal Transition, PLOS ONE 11 (2016) e0147816. 10.1371/journal.pone.0147816.

[35] N. Pandya Thakkar, B.M.V. Pereira, Y.T. Katakia, S.K. Ramakrishnan, S. Thakar, A. Sakhuja, G. Rajeev, S. Soorya, K. Thieme, S. Majumder, Elevated H3K4me3 Through MLL2-WDR82 upon Hyperglycemia Causes Jagged Ligand Dependent Notch Activation to Interplay with Differentiation State of Endothelial Cells, Front. Cell Dev. Biol. 10 (2022) 839109. 10.3389/fcell.2022.839109.

[36] X. Liang, S. Wu, Z. Geng, L. Liu, S. Zhang, S. Wang, Y. Zhang, Y. Huang, B. Zhang, LARP7 Suppresses Endothelial-to-Mesenchymal Transition by Coupling With TRIM28, Circ. Res. 129 (2021) 843–856. 10.1161/CIRCRESAHA.121.319590.

[37] G.J. Knott, C.S. Bond, A.H. Fox, The DBHS proteins SFPQ, NONO and PSPC1: a multipurpose molecular scaffold, Nucleic Acids Res. 44 (2016) 3989–4004. 10.1093/nar/gkw271.

[38] K.E. Lukong, M.-É. Huot, S. Richard, BRK phosphorylates PSF promoting its cytoplasmic localization and cell cycle arrest, Cell. Signal. 21 (2009) 1415–1422. 10.1016/j.cellsig.2009.04.008.

[39] K. Hirano, F. Erdödi, J.G. Patton, D.J. Hartshorne, Interaction of protein phosphatase type 1 with a splicing factor, FEBS Lett. 389 (1996) 191–194. 10.1016/0014-5793(96)00577-7.

[40] L. Liu, N. Xie, P. Rennie, J.R.G. Challis, M. Gleave, S.J. Lye, X. Dong, Consensus PP1 Binding Motifs Regulate Transcriptional Corepression and Alternative RNA Splicing Activities of the Steroid Receptor Coregulators, p54nrb and PSF, Mol. Endocrinol. 25 (2011) 1197–1210. 10.1210/me.2010-0517.

[41] B.A. Lewis, S.K. Das, R.K. Jha, D. Levens, Self-assembly of promoter DNA and RNA Pol II machinery into transcriptionally active biomolecular condensates, Sci. Adv. 9 (2023) eadi4565. 10.1126/sciadv.adi4565.

[42] J. Huang, M. Ringuet, A.E. Whitten, S. Caria, Y.W. Lim, R. Badhan, V. Anggono, M. Lee, Structural basis of the zinc-induced cytoplasmic aggregation of the RNA-binding protein SFPQ, Nucleic Acids Res. 48 (2020) 3356–3365. 10.1093/nar/gkaa076.

[43] T. Fukada, S. Yamasaki, K. Nishida, M. Murakami, T. Hirano, Zinc homeostasis and signaling in health and diseases: Zinc signaling, J. Biol. Inorg. Chem. JBIC Publ. Soc. Biol. Inorg. Chem. 16 (2011) 1123–1134. 10.1007/s00775-011-0797-4.

[44] M. Maywald, S.K. Meurer, R. Weiskirchen, L. Rink, Zinc supplementation augments TGF-β1-dependent regulatory T cell induction, Mol. Nutr. Food Res. 61 (2017). 10.1002/mnfr.201600493.

[45] J.J. Mack, T.S. Mosqueiro, B.J. Archer, W.M. Jones, H. Sunshine, G.C. Faas, A. Briot, R.L. Aragón, T. Su, M.C. Romay, A.I. McDonald, C.-H. Kuo, C.O. Lizama, T.F. Lane, A.C. Zovein, Y. Fang, E.J. Tarling, T.Q. De Aguiar Vallim, M. Navab, A.M. Fogelman, L.S. Bouchard, M.L. Iruela-Arispe, NOTCH1 is a mechanosensor in adult arteries, Nat. Commun. 8 (2017) 1620. 10.1038/s41467-017-01741-8.

[46] W.J. Polacheck, M.L. Kutys, J. Yang, J. Eyckmans, Y. Wu, H. Vasavada, K.K. Hirschi, C.S. Chen, A non-canonical Notch complex regulates adherens junctions and vascular barrier function, Nature 552 (2017) 258–262. 10.1038/nature24998.

[47] J. Zavadil, L. Cermak, N. Soto-Nieves, E.P. Böttinger, Integration of TGF-β/Smad and Jagged1/Notch signalling in epithelial-to-mesenchymal transition, EMBO J. 23 (2004) 1155– 1165. 10.1038/sj.emboj.7600069.

[48] M. Noseda, G. McLean, K. Niessen, L. Chang, I. Pollet, R. Montpetit, R. Shahidi, K. Dorovini-Zis, L. Li, B. Beckstead, R.E. Durand, P.A. Hoodless, A. Karsan, Notch Activation Results in Phenotypic and Functional Changes Consistent With Endothelial-to-Mesenchymal Transformation, Circ. Res. 94 (2004) 910–917. 10.1161/01.RES.0000124300.76171.C9.

[49] N. Rudini, A. Felici, C. Giampietro, M. Lampugnani, M. Corada, K. Swirsding, M. Garrè, S. Liebner, M. Letarte, P. ten Dijke, E. Dejana, VE-cadherin is a critical endothelial regulator of TGF-beta signalling, EMBO J. 27 (2008) 993–1004. 10.1038/emboj.2008.46.

