## Supplemental Figures for "Splicing factor proline- and glutamine-rich (SFPQ) protein causes transcriptional repression of SNAIL to counteract TGF-β signaling"

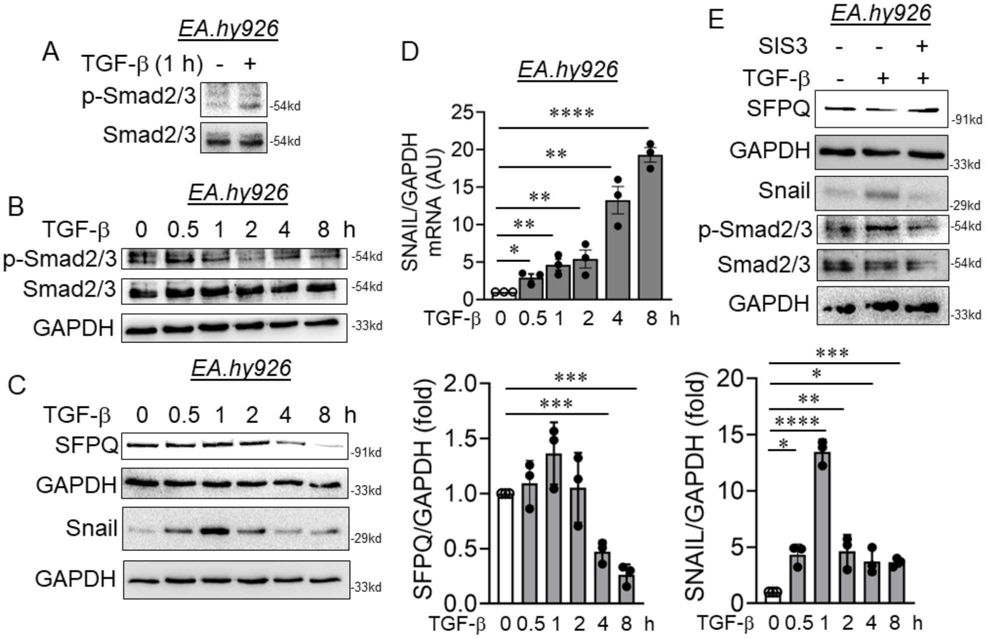


**Figure S1. TGF-β dependent regulation of SFPQ and SNAIL is dependent on Smad activation.** (A-C) EA.hy926 cells were treated with TGF-β (10 ng/mL) for different time points and the phosphorylation of Smad2/3 (A,B), SFPQ (C), and Snail (C) protein levels were detected in such cell lysates. (D) Transcript-level expression of *SNAIL* (n = 3) measured through RT-qPCR technique in EA.hy926 cells treated with TGF-β (10 ng/mL) for different time points. (E) EA.hy926 cells were pre-treated with pharmacological inhibitors of Smad phosphorylation followed by stimulating with TGF-β (10 ng/mL) for 4 h. Immunoblot analysis was carried out to detect the protein level of Snail, SFPQ, and phosphorylation of Smad2/3. Values represent the mean ± SD. * $p < 0.05$, ** $p < 0.01$, and **** $p<0.0001$ by one way ANOVA.

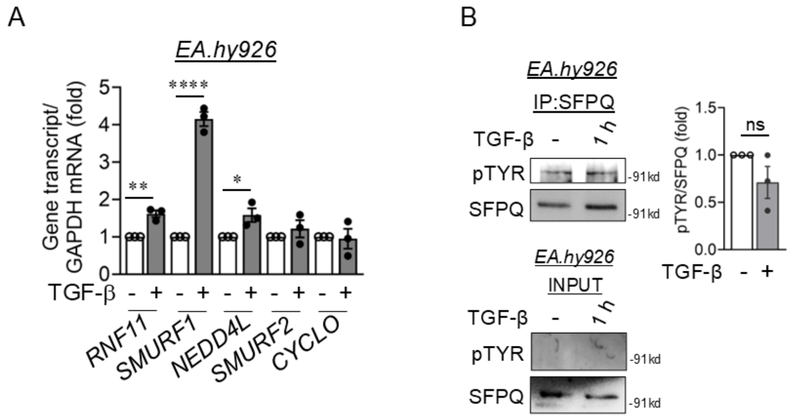


**Figure S2. TGF-β caused selective upregulation of E3 ubiquitin ligases and increase in tyrosine phosphorylation.** Transcript-level expression of *RNF11*, *SMURF1*, *NEDD4L*, *SMURF2*, and *CYCLO* (n = 3) measured through RT-qPCR technique in EA.hy926 cells treated with TGF-β (10 ng/mL) for 24 h. (B) EA.hy926 cells were treated with TGF-β for 1 hour and immunoprecipitated for endogenous SFPQ, fresh lysates were probed for phospho-tyrosine residues.

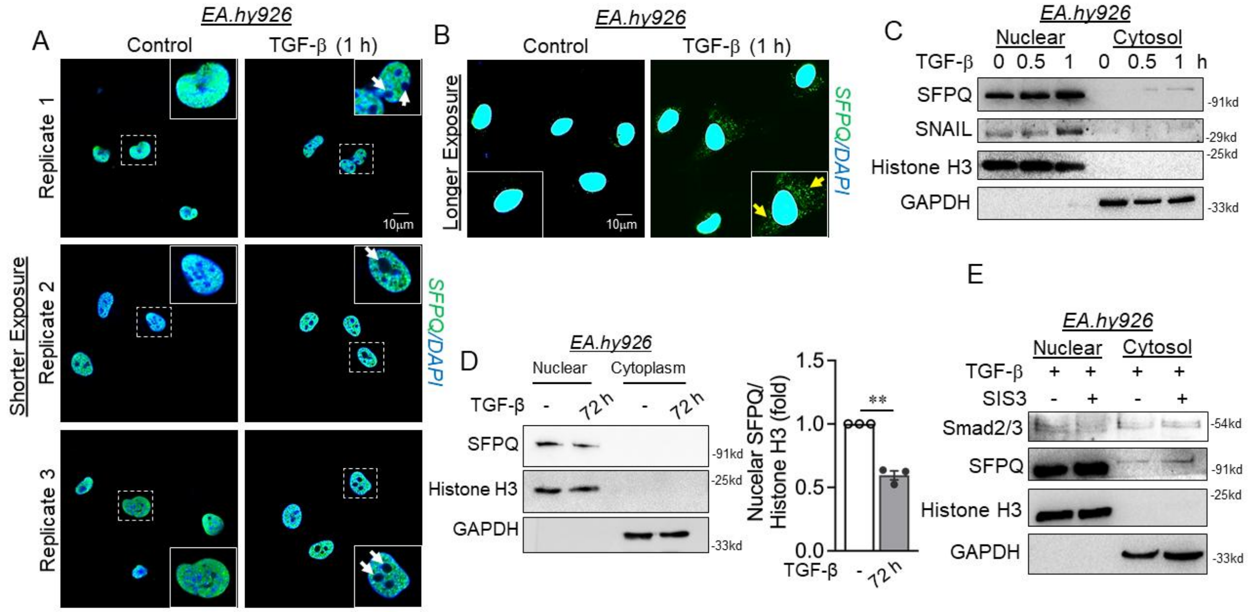


**Figure S3. TGF-β caused early cytosolic localization of SFPQ independent of Smad activation.** (A-B) Immunofluorescence staining of EA.hy926 cells exposed to TGF-β for 1 h and stained for SFPQ (Green). DAPI staining to visualize the nucleus is shown in blue. White arrow heads indicate the loss of SFPQ localization within the nucleus of the cells exposed to TGF-β.

Images were acquired in shorter (A) and longer exposure (B) settings to allow localization of SFPQ in cytosol of the cells. Yellow arrow heads indicate the presence of cytosolic SFPQ in the cells. (C-D) EA.hy926 cells were treated with TGF-β for 0.5, 1 h (C) or 72 h (D) and separated for cellular components, nuclear (10 μg) and cytoplasmic fraction (30 μg) were loaded and immunoblot for SFPQ protein, at late time interval 72 hours. GAPDH and histone H3 showed the purity of cytosolic and nuclear fractions respectively.(E) EA.hy926 cells were pre-treated with pharmacological inhibitors of Smad phosphorylation followed by stimulating with TGF-β (10 ng/mL) for 4 h. Cell lysates were separated for cellular components, nuclear (10 μg) and cytoplasmic fraction (30 μg) were loaded and immunoblot for SFPQ protein, at late time interval 72 hours. GAPDH and histone H3 showed the purity of cytosolic and nuclear fractions respectively. Values represent the mean ± SD. ** $p < 0.01$, by unpaired t test.

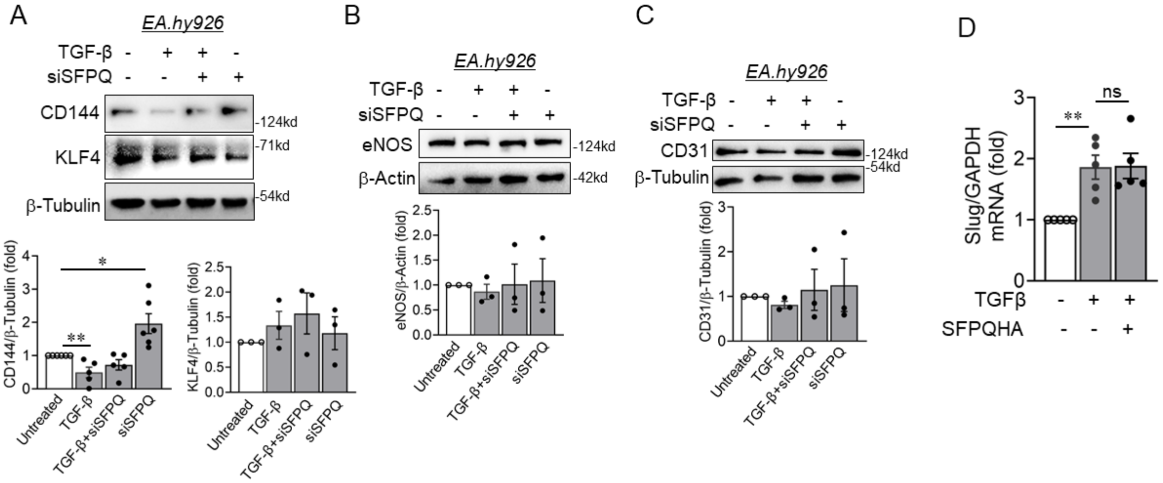


**Figure S4. Loss of SFPQ did not alter many of the endothelial specific genes except CD144 in TGF-β treatment conditions while TGF-β induced expression of transcription factor SLUG remained unchanged upon forced expression of SFPQ.** (A-C) Immunoblot analysis of EA.hy926 cell lysates transfected with siRNA against SFPQ were probed for CD144 (n = 5), KLF4 (n = 3), eNOS (B, n = 3), and CD31 (C, n=3). All analyzed data were normalized to the control treatment condition. (D) Transcript-level expression of *SLUG* (n = 5) measured through RT-qPCR technique in HEK293 cells expressing SFPQ exogenously and treated with TGF-β condition. Values represent the mean ± SD. * $p < 0.05$, and ** $p < 0.01$ by one way ANOVA.

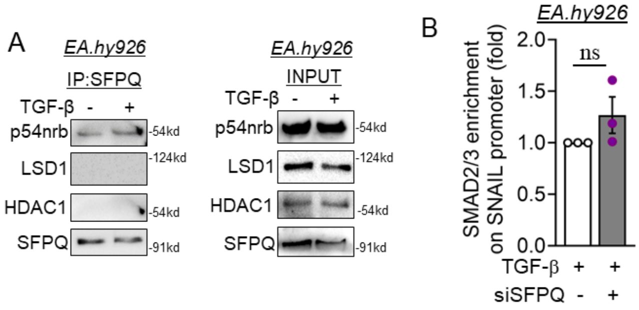


**Figure S5. SFPQ interaction with known binding partner p54nrb remained unchanged upon TGF-β stimulation while SFPQ did not associate with other nuclear proteins such as LSD1 and HDAC1.** (A) EA.hy926 cells were treated with TGF-β for 1 hour and immunoprecipitated for endogenous SFPQ, fresh lysates were probed for p54nrb, LSD1, and HDAC1 followed by comparing them with Input samples. (B) Chromatin occupancy study by Cut-and-run assay in EA.hy926 cell treated with TGF-β for 4 hours using Smad2/3 antibody followed by qPCR of eluent DNA with primers for the E-Box region of SNAIL gene promoter. All Ct values were normalized to the Ct value of control IgG.
