## Supplemental Table for "Splicing factor proline- and glutamine-rich (SFPQ) protein causes transcriptional repression of SNAIL to counteract TGF-β signaling"

**Supplementary Table 1: Sequences of the siRNA including scrambled and SFPQ siRNAs.**

| Gene specific siRNA | Target Sequence (5`-3`) |
|---|---|
| Custom designed SFPQ siRNA (Genecust, France) | GCACGUUUGAGUACGAAUAtt |
| | UAUUCGUACUCAAACGUGCtt |
| Silencer® SFPQ siRNA #AM16708 | GGCUAAGUAUUGCUUUCUAtt |
| | UAGAAAGCAAUACUUAGCCtc |
| Scramble siRNA (Genecust, France) | UUCUCCGAACGUGUCACGYtt |
| | ACGUGACACGUUCGGAGAAtt |

**Supplementary Table 2: Details of antibodies used in the study.**

| **Antibody name** | **Catalogue no.** |
|---|---|
| **Immunoblotting studies** | |
| Horse Anti-mouse IgG, HRP-linked Antibody (1:2000) | 7076 (CST) |
| Goat Anti-Rabbit IgG, HRP-linked Antibody (1:2000) | 7074 (CST) |
| Histone H3 (96C10) Mouse monoclonal antibody | 3638 (CST) |
| VE-Cadherin (D87F2) XP(R) Rabbit monoclonal antibody | 2500 (CST) |
| Snail (C15D3) Rabbit monoclonal antibody | 3879 (CST) |
| GAPDH (D16H11) XP® Rabbit monoclonal antibody | 5174 (CST) |
| β-Tubulin (D3U1W) Mouse monoclonal antibody | 86298 (CST) |
| KLF4 Rabbit Polyclonal antibody | 4038 (CST) |

| | |
|---|---|
| Ubiquitin (P4D1) Mouse monoclonal antibody | 3936 (CST) |
| HA-Tag (C29F4) Rabbit monoclonal antibody | 3724 (CST) |
| SMAD2/SMAD3 Rabbit Polyclonal antibody | A7536 (AbClonal) |
| Smad4 Rabbit monoclonal antibody | A19116 (AbClonal) |
| SFPQ Rabbit monoclonal antibody | A3494 (AbClonal) |
| Notch1(Cleaved) Rabbit Polyclonal antibody | A16673 (AbClonal) |
| Anti-PSF Mouse monoclonal antibody | P2860 (Sigma Aldrich) |
| Pan Phospho-Tyrosine Rabbit Polyclonal antibody | AP0905 (AbClonal) |
| P54nrb Rabbit Polyclonal antibody | A300-587A (Bethyl Lab) |
| PSPC1 Rabbit Polyclonal antibody | A303-206A (Bethyl Lab) |
| SQSTM1/p62 Rabbit monoclonal antibody | 39749 (CST) |
| LC3A/B Rabbit monoclonal antibody | 12741 (CST) |
| eNOS Rabbit monoclonal antibody | 32027 (CST) |
| CD31 (PECAM-1) Mouse monoclonal antibody | 3528 (CST) |
| LSD1 Mouse monoclonal antibody | 4218 (CST) |
| HDAC1 Mouse monoclonal antibody | 5356 (CST) |
| JG12 Mouse monoclonal antibody | BMS1104 (Thermo Fisher) |
| SFPQ Rabbit monoclonal antibody | Ab177149 (Abcam) |

**Supplementary Table 3: Sequences of the primer used to measure the transcript level of different genes using qPCR.**

| Gene name | Forward sequence | Reverse sequence |
|---|---|---|
| *SFPQ* | *5'AGAGGAAAGTTACAGCCGAATGG3'* | *5'CATAACCTATGCCACCACCACCTCG3'* |
| *Snai1* | *5'CGAGTGGTTCTTCTGCGCTA3'* | *5'GGGCTGCTGGAAGGTAAACT3'* |
| | *5'GAGCCCAGGCAGCTATTTCA3'* | *5`TGGGAGACACATCGGTCAGA3'* |
| *Slug* | *5'AACAGTATGTGCCTTGGGGG3'* | *5'AAAAGGCACTTGGAAGGGGT3'* |
| *Rnf11* | *5' CACCAATCCCTCAGCATCTT 3'* | *5' TTCCATCCTCTGCATCACTTC 3'* |
| *Nedd4l* | *5' TGAATCTGGCTGCTGGTGTA 3'* | *5' TCTAGCACACAAGAGGCACA 3'* |
| *Smurf1* | *5' TCTTTGAGGAGTCTTACCGC 3'* | *5' CACACCACCGTAATCCAAAC 3'* |
| *Smurf2* | *5' GACAGTCTTCAGATCCCAGG 3'* | *5' TGCGTTGTCCTCTGTTCATA 3'* |
| *Cyclophilin A* | *5' ATGGTCAACCCCACCGTG 3'* | *5' TGCAATCCAGCTAGGCATG 3'* |

**Supplementary Table 4: Sequences of the SNAIL gene promoter/coding region primers used to measure the enrichment of SFPQ.**

| Gene Promotor | Forward sequence | Reverse sequence |
|---|---|---|
| *hSNAIL TSS (ref point)* | *5'GGAGTACTTAAGGGAGTTGGCGG3'* | *5'GAACCACTCGCTAGGCCGT 3'* |
| *hSNAIL TSS #2* | *5'GGCCTAGCGAGTGGTTCTTC3'* | *5'CCTCCAACGCACCTGGATTAG3'* |
| *hSNAIL plus 260bp* | *5'GCGAGCTGCAGGACTCTAAT3'* | *5'CCCGCAATGGTCCACAAAAC3'* |

| | | |
|---|---|---|
| *hSNAIL plus 558bp* | *5'TGTTTTGTGGACCATTGCGG3'* | *5'GAGGCCAGAACCCCATCATC3'* |
| *hSNAIL plus 654bp* | *5'ATGTTTTGTGGACCATTGCGG3'* | *5'CCCAGGTCACCCAATCTGAG3'* |
| *hSNAIL plus 770bp* | *5'GACTCAGATTGGGTGACCTGG3'* | *5'ACTCAATCAACAAACATGAGCCC3'* |
| *hSNAIL minus 654bp* | *5'CGGGAGAGGCTCTGAGTGTT3'* | *5` CTAGCCAAGAGCACCCGTTC 3`* |
